# Saturating hepatic clearance drives elevated cfDNA and fragment shortening in cancer

**DOI:** 10.64898/2026.03.04.709433

**Authors:** Thomas Rachman, William A. LaFramboise, Phillip H. Gallo, Patricia Petrosko, Daisong Liu, Rahul Kumar, Marija Balic, Steffi Oesterreich, Julia Foldi, Adrian V. Lee, Patrick L. Wagner, David L. Bartlett, Russell Schwartz, Oana Carja

## Abstract

Liquid biopsy studies consistently report both elevated circulating cell-free DNA (cfDNA) concentrations and shortened fragment lengths in cancer. These features are often attributed to tumor-specific processes, despite tumor-derived cfDNA frequently constituting less than 1% of the total. Here, we consider an alternative explanation: Saturation of cfDNA clearance, which prolongs cfDNA circulation time, increases exposure to plasma nucleases and is expected to produce similar fragmentomic signatures independent of tumor burden. By combining a mechanistic model of cfDNA fragmentation with analyses of two independent cancer patient cohorts, and publicly available clearance-perturbation experiments, we demonstrate that elevated cfDNA levels are accompanied by a characteristic leftward shift in fragment length distributions consistent with impaired hepatic clearance. This fragmentation signature becomes more pronounced at higher cfDNA concentrations, is independent of circulating tumor DNA (ctDNA) fraction, is reproducible under experimentally reduced clearance, and is independently prognostic of patient survival. Together, these results identify saturating clearance as a central determinant of cfDNA abundance and fragment length, re-framing cancer-associated fragmentomic patterns as systemic consequences of clearance dynamics rather than tumor burden alone. More broadly, they highlight the value of mechanistic modeling of clearance processes in extracting clinically meaningful signals from cfDNA fragmentation data.

## Introduction

Liquid biopsies have emerged as a powerful, noninvasive approach for cancer detection and disease monitoring, with circulating cell-free DNA (cfDNA) serving as a central biomarker [1, 2, 3, 4]. Both elevated cfDNA concentrations and altered fragment length distributions are consistently associated with disease progression and poor clinical outcomes, motivating their recent widespread incorporation into machine-learning–based diagnostic and prognostic models [5, 6, 7, 8, 9].

A key observation underlying these approaches is that tumor-derived fragments are, on average, shorter than those originating from healthy cells [10, 11, 12]. As a result, the global cfDNA shortening in cancer patients is commonly attributed to tumor-specific biological processes, such as altered nucleosome organization or cancer-associated modes of cell death [6, 13, 14, 7]. However, global reduction in cfDNA fragment length is difficult to reconcile with the typically low tumor fraction in plasma. Despite markedly elevated total cfDNA concentrations, tumor-derived cfDNA often constitutes less than 1% of the total, with hematopoetic cells contributing the majority of circulating fragments [15, 16, 17, 18]. Tumor cell turnover alone is therefore unlikely to completely account for both the increased cfDNA abundance and the systematic shortening of fragment lengths observed in cancer patients.

An alternative, non–tumor-specific explanation may lie in the saturation of cfDNA clearance pathways. If cfDNA removal becomes capacity-limited, even modest increases in DNA shedding could lead to disproportionate increases in steady-state cfDNA levels. The resulting prolonged circulation times would, in turn, increase exposure to circulating nucleases, promoting progressive fragment shortening. Clearance saturation could therefore reconcile elevated cfDNA concentrations with reduced fragment lengths, even in the presence of low tumor fractions.

Here, we ask whether fragment shortening and elevated cfDNA levels in cancer patients primarily reflect tumor-derived DNA or, instead, arise from systemic clearance saturation and prolonged circulation time. CfDNA is cleared predominantly in the liver by Kupffer cells, which are likely subject to a finite maximal clearance capacity [19, 20]. Consistent with this hypothesis, recent fragmentomics analyses of sepsis patients report both elevated cfDNA concentrations and shorter fragments, attributed to impaired hepatic clearance and increased circulation times [21]. Moreover, methylation-based analyses in early-stage cancer suggest cfDNA dynamics are better described by saturating rather than linear clearance models [22]. Together, these observations motivate a systematic investigation of clearance saturation as a driver of cfDNA fragmentation in cancer.

We address this question by combining a mathematical model of cfDNA fragmentation with analyses of public experimental data that perturb hepatic and enzymatic clearance in mice [23], alongside two independent human cancer cohorts. Our work aims to unify cfDNA shedding, clearance saturation, circulation time, and nuclease-driven fragmentation into a single framework. We favor this mechanistic approach over purely ML-based strategies because, while these approaches excel at prediction, mechanistic models enable direct interrogation of the biological processes shaping fragment length distributions. Existing models either focus on nucleosome occupancy without perturbing fragmentation [24] or describe kilobase-scale fragmentation while ignoring nucleosome protection [25]. In order to make use of widely available short-read fragmentomics data, our model jointly examines fragmentation and clearance at the mononucleosome scale.

We show that saturation of hepatic clearance accounts for the concurrent elevation in cfDNA concentrations and reduction in fragment lengths observed in cancer patients. Furthermore, we demonstrate that ctDNA content alone cannot explain the relationship between cfDNA concentration and fragment length. Reduced clearance is associated with a shift toward shorter modal fragment lengths, which we identify as a robust fragmentomic feature with independent prognostic significance. Collectively, our work reframes cfDNA fragment length distributions, particularly the structure of the mononucleosome peak, as emergent properties of fundamental physical and biological constraints. This reinterpretation has broad implications for fragmentomics-based inference and clinical decision support in cancer and other diseases characterized by high cfDNA burden.

## Results

### Mathematical model of cfDNA yield and fragmentation under saturating clearance

#### cfDNA yield dynamics

To investigate whether cfDNA fragment shortening and elevated cfDNA levels in cancer patients primarily reflect tumor burden or can instead arise from systemic clearance saturation and prolonged circulation time, we first compare a linear clearance process with a saturating clearance mechanism governed by Michaelis–Menten kinetics. We assume that cfDNA concentration is at quasi–steady state at the time of blood draw. This assumption is supported by the short half-life of cfDNA in plasma, the rapid activity of DNases observed in vitro relative to typical rates of cell division and death, and its use in prior models of cfDNA kinetics [26, 27, 25, 28, 22, 29]. We define the shedding rate *s* to be the total inflow of cfDNA molecules per unit time, proportionate to the total volume of shedding cells, and the hepatic clearance rate *k* to be the rate of fragment exit per unit time.

Under linear clearance, total cfDNA concentration scales proportionally with the shedding rate *s*,

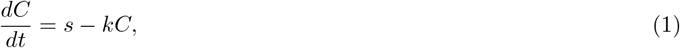

such that the equilibrium concentration is determined by the ratio of shedding to clearance,

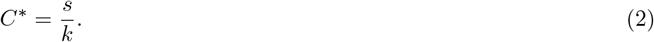

We contrast this with a saturating clearance model in which cfDNA removal follows Michaelis–Menten kinetics, where *V*_max_ denotes the maximal clearance capacity and *K*_*m*_ the half-saturation constant,

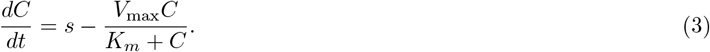

Unlike the linear case, this regime allows cfDNA concentration and circulation time to increase disproportionately as shedding approaches the clearance capacity. At steady state, the cfDNA concentration is given by

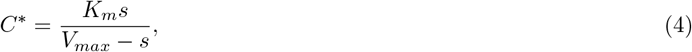

and the effective per-molecule clearance rate becomes (*V*_max_ − *s*)*/K*_*m*_, which decreases toward zero as the shedding rate *s* approaches *V*_max_.

In **Figure 1A**, we show the effect of increasing the shedding rate on the equilibrium cfDNA concentration for both models. When increased shedding is driven by tumor growth, these dynamics predict disproportionately elevated cfDNA levels for large tumors (**Figure 1B**).

**Figure 1:**
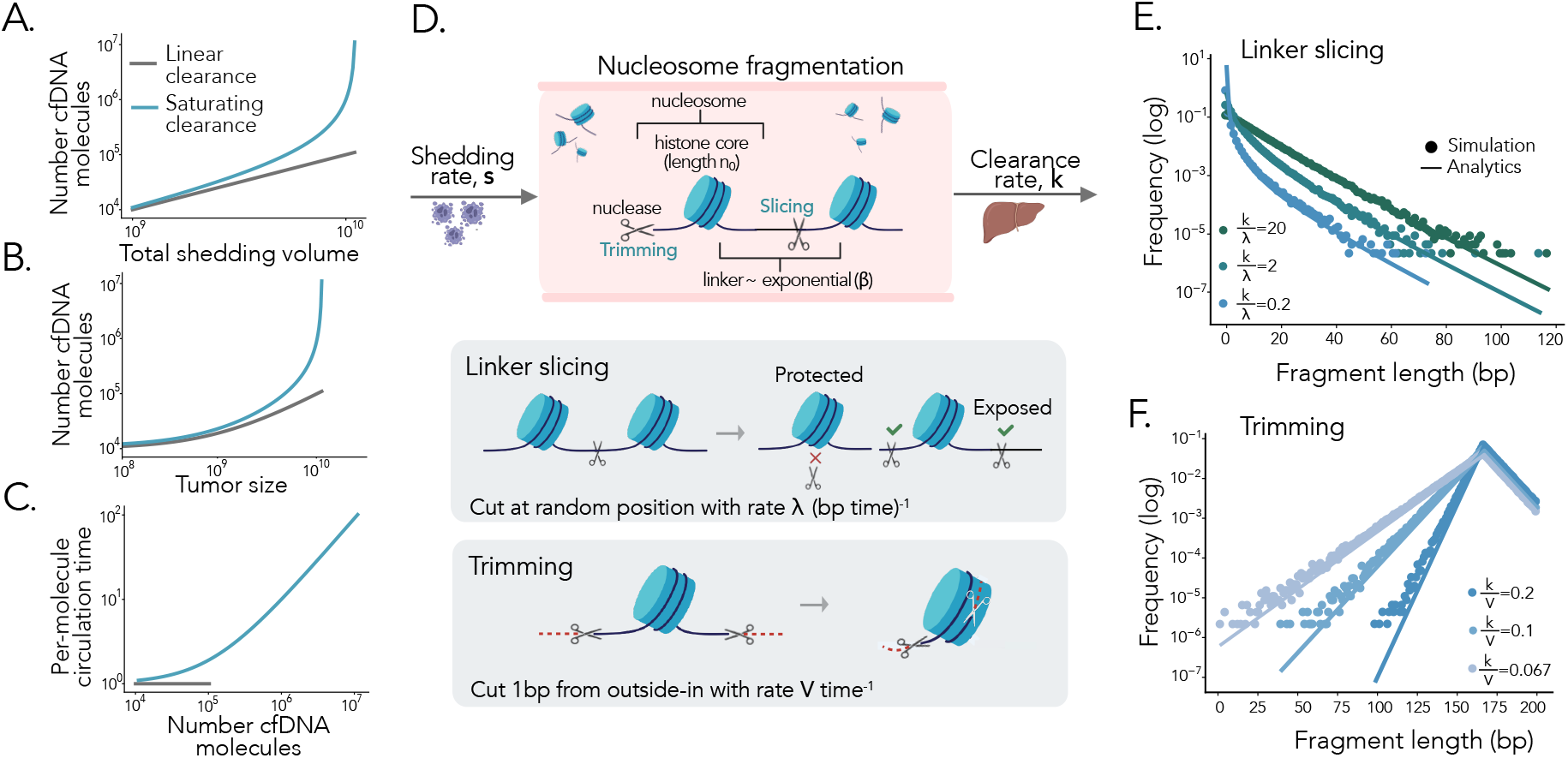
A model of saturating clearance and fragmentation. **A:** Example trajectory of cfDNA amount when clearance rate is constant (gray) versus saturating (blue), as a function of total shedding volume. The lines show the equilibria in equations (2) and (4). **B:** Example trajectory of cfDNA amount as a function of tumor size, assuming a constant non-tumor shedding source and a growing tumor source. **C:** Circulation time (the inverse of the effective clearance rate) plotted for the same trajectory. As circulation time increases, fragments naturally receive more cuts. **D:** Illustration of the fragmentation model. Mononucleosomal DNA fragments consists of a protected region of 167bp, controlled by the nucleosome wrap of length *n*_0_, and an exposed region, with length drawn from an exponential distribution with rate *β*, plus some minimal exposed value *l*_0_, initially set to 0. Fragments are shed into the bloodstream with rate *s*. Upon entering circulation, they are cut via slicing and trimming. Slicing occurs on exposed linker regions at a rate *λ* bp^−1·^time^−1^. The slice rate is scaled by the linker length, so that fragments with more exposed linker are cut more rapidly. Trimming occurs at a constant rate and removes one basepair at a rate *v* time^−1^. Fragments are uniformly sampled for clearance with rate *k*, which effectively captures both saturating and linear clearance models at equilibrium. **E:** Analytical prediction for the stationary distribution of fragment length for the slicing process on an exponentially-distributed linker fragment overlaid with results of exact stochastic simulation for a range of relative fragmentation rates (*s* = 10^8^, *k* = 200, *β* = 0.1, *λ* = 10, 10^2^, 10^3^). **F:** Analytics and simulations for the trimming process on a complete mononucleosome with constant region *n*_0_ flanked by an exponential linker (*n*_0_ = 167, *s* = 10^8^, *k* = 200, *β* = 0.1, *λ* = 10, *v* = 1000, 2000, 3000).

#### Fragment length dynamics

Under a saturating clearance model, the per-molecule circulation time increases, leading to prolonged persistence of individual molecules in plasma (**Figure 1C**). We hypothesize that this extended residence time results in greater cumulative fragmentation at higher concentrations. To evaluate the impact of increased fragmentation, we model nucleosome-bound DNA entering circulation, undergoing fragmentation and clearing. This allows us to characterize the resulting equilibrium distribution of cfDNA fragment lengths. We focus on fragments between 100 and 200 bp, which dominate short-read sequencing libraries prepared from double-stranded DNA [11, 30].

We initialize fragments as nucleosome-bound DNA composed of a protected core and an exposed linker region (**Figure 1D**). Fragments in this size range arise when nucleases *slice* exposed linker DNA between adjacent nucleosomes. We model cuts as occurring uniformly at random along the exposed region, assumning a constant slice rate *λ* per bp·time and an initial linker length that is exponentially distributed with rate *β* [24]. This generates a power-law decay in fragment length distributions, reflecting repeated cleavage of exposed linker DNA. Specifically, exposed regions initially exhibit exponential decay, with progressively steeper right-tail behavior as fragmentation increases relative to clearance. For a fragment of length *x >* 0 the stationary distribution *f*_slice_(*x*) is given by (see **Supplementary Material Section 1.1.3** for derivations):

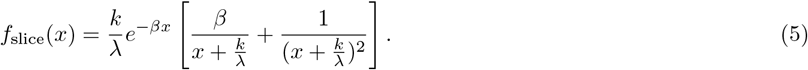

Slicing alone, however, cannot account for the pronounced mononucleosomal peak commonly observed near 167 bp and the gradual increase in frequency of fragments before this peak [7]. To capture this feature, we additionally allow fragments to be shortened from the ends inward, a process we term *trimming*. In circulating plasma, nucleases preferentially cleave histone-bound DNA at outward-facing minor grooves, producing characteristic oscillations in fragment length at 10.4 bp intervals [31, 7]. Rather than explicitly model this fine-scale periodicity, we focus on the net effect of progressive end erosion up to the nucleosome boundary. We approximate trimming as a constant, deterministic erosion of fragment ends at rate *v* (bp·time^−1^), applied to all fragments regardless of nucleosome protection.

Separating cases based on whether trimming has reached the nucleosome boundary, which occurs at *x* = *n*_0_, yields the stationary distribution *f*_trim_ (see **Supplementary Material Section 1.1.4** for derivations):

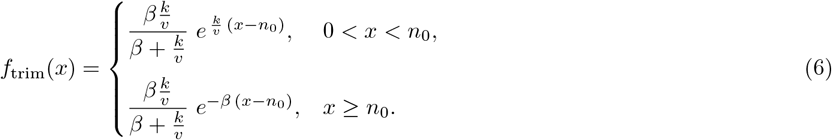

Exact stochastic simulations of both cutting processes closely match these analytical predictions (**Figure 1E–F**). Notably, the left tail of the fragment length distribution is governed entirely by the ratio of clearance to trimming, while the right tail depends on both the ratio of clearance to slicing and the initial linker length distribution. Despite these mechanistic differences, both models yield the same qualitative conclusion: the shape of the stationary fragment length distribution is controlled by the ratio of fragmentation to clearance, rather than by their absolute rates.

### Comparison to clearance perturbation experiments in mice

To examine the combined effects of fragmentation on mononucleosome-length cfDNA and to compare model predictions with fragmentation data, we integrate the two fragmentation mechanisms, slicing and trimming, into a single stochastic simulation. The nucleosome-bound DNA only undergoes trimming, while the exposed linker DNA is subject to both forms of cutting. We first generate a baseline distribution using arbitrary shedding and clearance parameters, protecting 167 bp of DNA from slicing (*n*_0_ = 167), which produces a modal fragment length at 167 bp. We then independently increase the two fragmentation rates, leading to the predicted narrowing of the right tail and broadening of the left tail of the fragment length distribution for slicing and trimming respectively, consistent with our analytical results (**Figure 2A–B**). Using this model, we assess how changes in fragmentation rates reshape the steady-state distribution of cfDNA fragment lengths.

**Figure 2:**
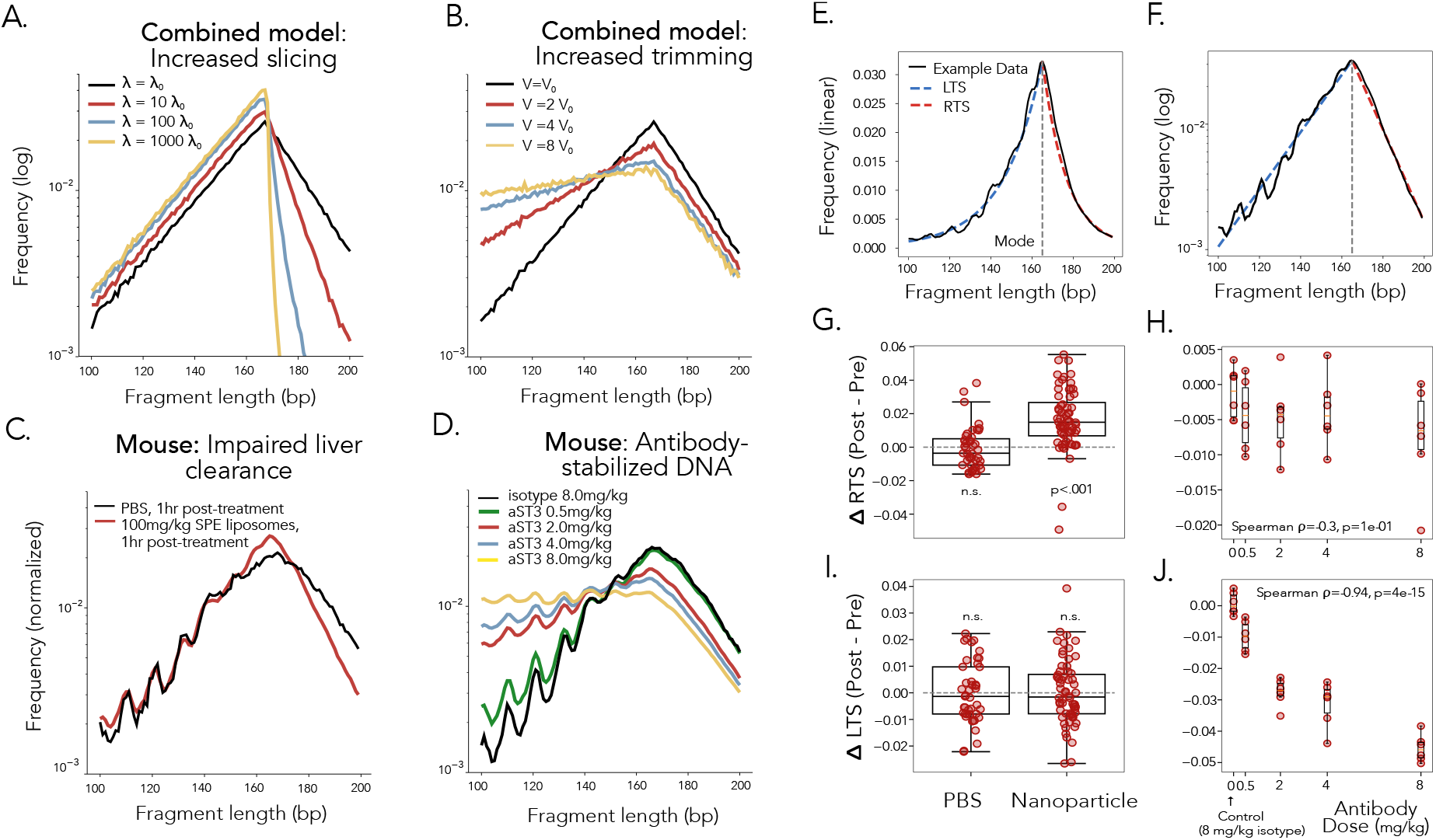
Fragmentation patterns in clearance-impaired mice recapitulate modeling results. **A:** Combined model with baseline parameters (*s* = 1*e*8, *λ* = 10, *v* = 4651, *k* = 200, *β* = 0.043, *n*_0_ = 167, *l*_0_ = 0), showing the effect of increasing *λ*. **B:** Combined model with baseline identical to panel A and increasing *v*. **C:** Aggregate length distribution for the Kupffer cell-blocking nanoparticle and PBS-treated control mice one hour after treatment. **D:** Aggregate length distribution for mice treated with 0.5-8mg of nucleosome-binding antibody aST3 and 8mg of inactive control (isotype) two hours after treatment. **E-F:** Absolute left tail slope (LTS) and right tail slope (RTS) fits in linear and log-frequency space to a single mouse sample. **G:** Change in RTS in paired samples before and 1 hour after PBS and nanoparticle administration. **H:** Change in RTS for paired samples before and 2 hours after antibody administration, plotted by dosage. The control dose of 8mg isotype antibody is shown at 0mg/kg. **I:** Change in LTS for control and nanoparticle treatments. **J:** Change in LTS for antibody treatments.

To compare model predictions with experimental data, we first analyze cfDNA fragmentation profiles from clearance perturbation experiments in mice, previously reported by Martin-Alonso et al. [23], hereafter referred to as the MA dataset. In these experiments, cfDNA clearance is transiently reduced using two distinct interventions: a lipid nanoparticle that blocks Kupffer cell–mediated hepatic clearance and a nucleosome-binding antibody designed to disrupt enzymatic degradation. Relative to their respective controls (phenyl-buffered saline and an inactive isotype antibody, shown as black lines in **Figure 2C–D**), these interventions produce distinct alterations in the mononucleosome region of the fragment length distribution. Nanoparticle-mediated inhibition of hepatic clearance yields a narrower right tail beyond the mononucleosomal peak, consistent with a model of increased degradation of exposed linker DNA (**Figure 2A, C**), whereas antibody treatment increases the abundance of fragments shorter than the modal length, resulting in a pronounced broadening of the left tail, consistent with a model of increased rates of trimming (**Figure 2B, D**). No differences were observed between pre-treatment groups (**Supplemental Figure S1)**. More generally, the results demonstrate that interventions acting on mechanisms responsible for distinct steps in DNA clearance modulate distinct parameters of the model that can be fit to experimentally observed fragment length distributions.

#### Quantifying changes in fragment length distributions

To quantify the observed changes in mononucleosomal cfDNA fragment profiles across experiments, we leverage two key results from our model. First, the model predicts that the ratio of clearance to trimming determines the slope of the left tail of the fragment length distribution. We estimate this empirically by computing the absolute slope of the best-fit line in log-frequency space from 100 bp up to the modal fragment length within the 100–200 bp range; we refer to this metric as the *left-tail slope* (LTS) (see **Supplementary Methods, Supplementary Figure S2A**).

Second, we compute the absolute slope from the modal length to 200 bp, which we term the *right-tail slope* (RTS). When clearance is high relative to slicing, the RTS is determined solely by the initial linker length distribution. It then increases monotonically as the slice rate increases relative to clearance, so that high linker fragmentation produces a steep right tail.

While it is possible to fit *f*_*slice*_ directly to the combined model, parameter inference is impacted by *v*, while RTS is not affected by changes to *v* and is therefore a more robust signature of 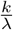 (**Supplementary Figure S2B-C**). Together, the LTS and RTS provide independent quantitative measures of the two fragmentation mechanisms relative to the global clearance rate. Because both quantities are computed in log-frequency space, they are robust to differences in normalization and sequencing depth.

Representative fits for the left-tail slope (LTS) and right-tail slope (RTS) in the MA dataset are shown in **Figure 2E–F**. In **Supplementary Figure S3**, we report the Pearson correlation coefficients for the LTS and RTS fits across all treatments and time points, which generally exceed 0.9, indicating robust linear behavior despite the expected periodicity in the left tail. To corroborate the differences observed in the aggregate cfDNA fragment length distributions (**Figure 2E–F**), we compute LTS and RTS values for each sample and analyze paired differences within individual mice before and after treatment. Nanoparticle-treated mice show a significant increase in RTS one hour post-treatment (Wilcoxon test, *p <* 0.001, *n* = 64), whereas control animals show no significant change (*p* = 0.10, *n* = 43) (**Figure 2G**). In contrast, LTS does not change significantly in either nanoparticle-treated or control, PBS-treated mice (**Figure 2I**).

Antibody-treated mice, however, exhibit a significant decrease in LTS two hours after treatment, while the control group shows no significant change (Wilcoxon test, *p <* 0.001 and *p* = 0.8, respectively)(**Figure 2J**). Moreover, the magnitude of the LTS decrease is strongly dose dependent (Spearman *ρ* = −0.95, *p <* 0.001). In this group, RTS does not change significantly in either the treatment or control groups (**Figure 2H**). These results demonstrate that we can quantify the effect of impaired clearance on an individual subject using log-linear slopes in the 100-200bp range of the fragment profile.

### Tumor fraction does not explain observed fragmentation signatures

Because circulating tumor DNA (ctDNA) fragments are typically shorter and more variable in length than non-tumor cfDNA, elevated tumor fraction could, in principle, account for both increased cfDNA abundance and reduced fragment length. However, in the original analysis of the MA dataset, tumor fractions remained stable or decreased slightly following clearance reduction, and were generally below 1% in the treatment group. These low tumor fractions are therefore unlikely to substantially influence fragment length distributions in this setting. Nevertheless, at sufficiently high levels, tumor-derived cfDNA would be expected to measurably alter fragment length profiles.

We extend our model to explicitly account for this possibility and, to determine whether tumor fraction alone can drive the fragment length changes predicted by our model, we evaluate cfDNA fragment length distributions in two independent clinical cohorts, spanning a wide range of tumor burdens. *Dataset I* is a pan-cancer cohort (*n* = 696) spanning 25 organ sites and all disease stages, comprising samples from patients enrolled in the Allegheny Health Network (AHN) Moonshot program. A clinical analysis of a subset of this cohort has been reported previously [32], with additional cohort details provided in **Supplementary Tables S1** and **S2**. All cfDNA samples in Dataset I are sequenced using the TruSite Oncology 500 gene panel (TSO500). *Dataset II* consists of samples from patients with HR+HER2-metastatic invasive lobular breast cancer (ILC) with *n* = 95 samples from 31 individuals included in the University of Pittsburgh Breast Disease Research Repository. All samples in Dataset II are sequenced via ultra-low-pass whole-genome sequencing. A breakdown of cfDNA and tumor fraction by patient is given in **Supplementary Table S3**.

To model the effects of tumor-derived fragmentation, we allow “protected” cfDNA to undergo fragmentation within the 143–167 bp range, such that at high slicing rates the modal fragment length converges to 143 bp, a widely observed mode for tumor-derived fragments [33]. Mixing this shorter-fragment distribution with the baseline cfDNA distribution yields a bimodal fragment length profile (**Figure 3A**).

**Figure 3:**
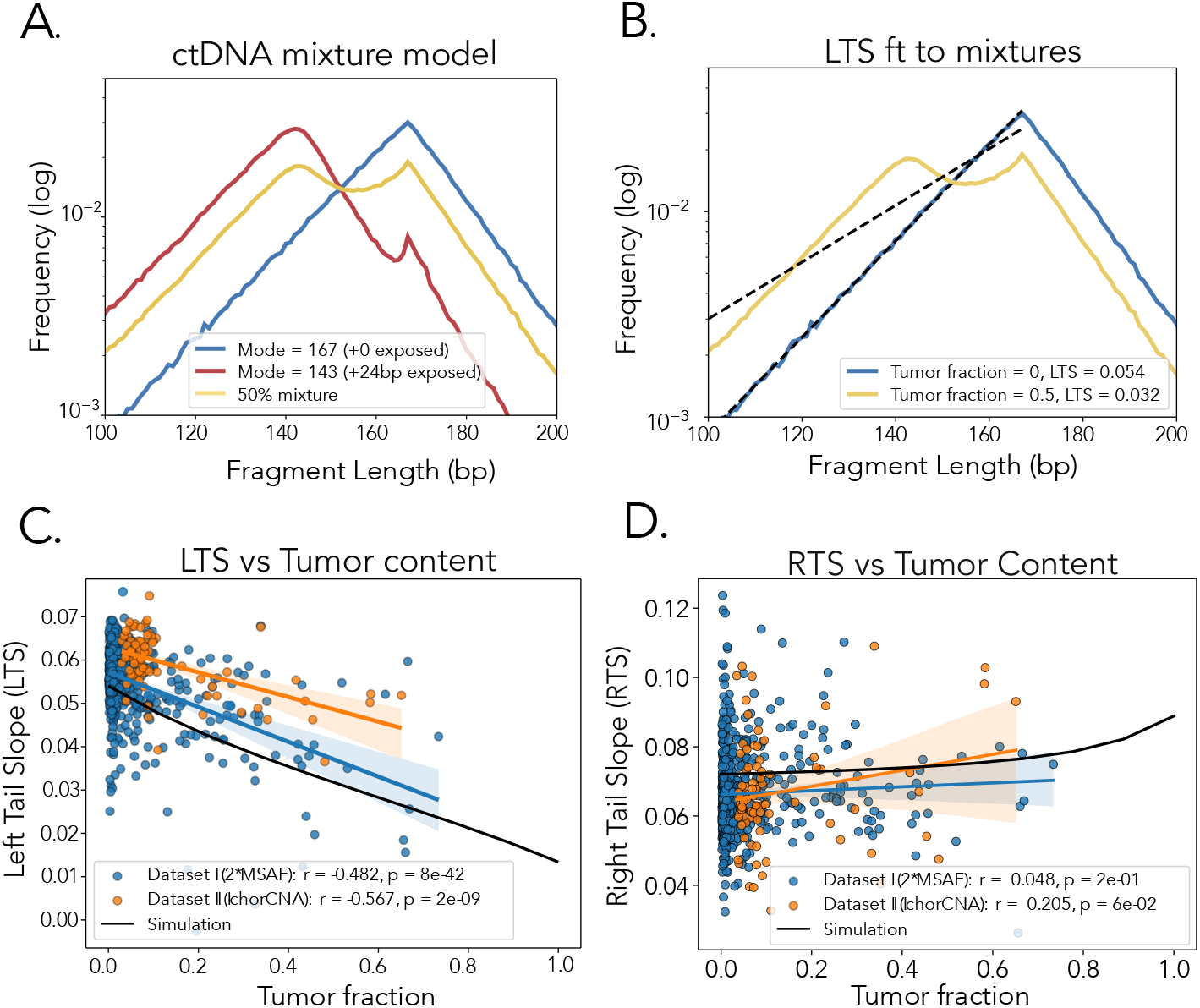
CtDNA impacts fragment profiles, but does not account for global shortening. **A:** Simulation of highly fragmented cfDNA created by increasing *λ* with respect to the baseline and allowing up to 24bp of the nucleosome wrap to degrade via slicing, meant to resemble ctDNA, mixed with a baseline distribution (*s* = 10^8^, *β* =.059, 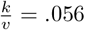, *λ* = 10, *n*_0_ = 167, *l*_0_ = 0). Parameters for the fragmented DNA are the same as the baseline but with *λ* = 100, *n*_0_ = 143, *l*_0_ = 24. **B:** Line of best fit for computing the LTS from the original fragment mode at 167bp, before and after mixing with 50% ctDNA. **C:** The LTS computed for the scenario in B for mixtures ranging from 0 to 100% ctDNA, showing the expected downward trajectory for the LTS value plotted with the LTS values and Pearson correlations for Dataset I (multistage, pan-cancer, n=696) and Dataset II (stage IV breast cancer, n=95). Samples were included if the tumor fraction was within the detection limit for each method (0.1%for MSAF, 3% for IchorCNA). **D:** Correlations between RTS and tumor fraction for the same samples, along with the computed RTS for the same simulated mixtures.

Let us first focus on the effect on the LTS. When LTS is calculated from 100 bp to the baseline mode at 167 bp, the additional tumor-derived mode distorts the linear fit to the left tail, producing a decrease in LTS relative to the 0% mixture (**Figure 3B**). By performing this fit for a range of mixtures, we can simulate the expected effect of tumor fraction on LTS.

We calibrate the combined model by setting 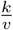 to the median LTS among samples with an MSAF (maximum somatic allele frequency) below 0.1% and *β* to the median RTS among the bottom decile by cfDNA concentration, which are 0.056 and.059 respectively. In **Figure 3C**, we show excellent agreement between our simulation and the two patient datasets, where LTS is negatively associated with tumor fraction. In contrast, we observe no correlation between the RTS and tumor fraction in either cancer cohort, along with very little change to the simulated RTS for the range of tumor fractions in the patient data (**Figure 3D**). Together, these results indicate that while sufficiently high tumor fractions alter cfDNA fragment length profiles, this effect is constrained to the left tail and unlikely to influence signatures of hepatic clearance in the right tail.

### cfDNA fragmentation in independent cancer cohorts reflects hepatic clearance

To determine whether clearance-associated fragmentation patterns extend to human data, we compared cfDNA fragment length distributions in mouse and human cohorts as a function of cfDNA concentration. Stratifying samples in both cohorts by cfDNA decile reveals systematic shifts in fragment length distributions that mirror the effects of enhanced linker fragmentation and nanoparticle-mediated clearance inhibition observed in mice (**Figure 4A–B**). Despite differences in clinical composition and sequencing methodology, the RTS exhibits a nearly identical relationship with cfDNA concentration across both human cohorts (**Figure 4C**). Consistent with this pattern, the RTS similarly increases with cfDNA concentration in mice treated with 100 mg/kg lipid nanoparticles, supporting a model in which progressive hepatic clearance saturation leads to increased fragmentation (**Figure 4C**). In Dataset I, patients with primary or metastatic liver involvement display elevated cfDNA concentrations and RTS values, both overall and within stage III–IV (**Figure 4D-E**). Although liver involvement is also associated with higher tumor fraction, its relationship to fragment length mirrors that observed previously in **Figure 3C-D** (see also **Supplementary Figure S4A-C**). Moreover, cfDNA concentration and RTS do not differ significantly between primary and metastatic liver involvement, suggesting that the presence of cancer within the liver broadly impacts clearance dynamics (**Supplementary Figure S4D-E**).

**Figure 4:**
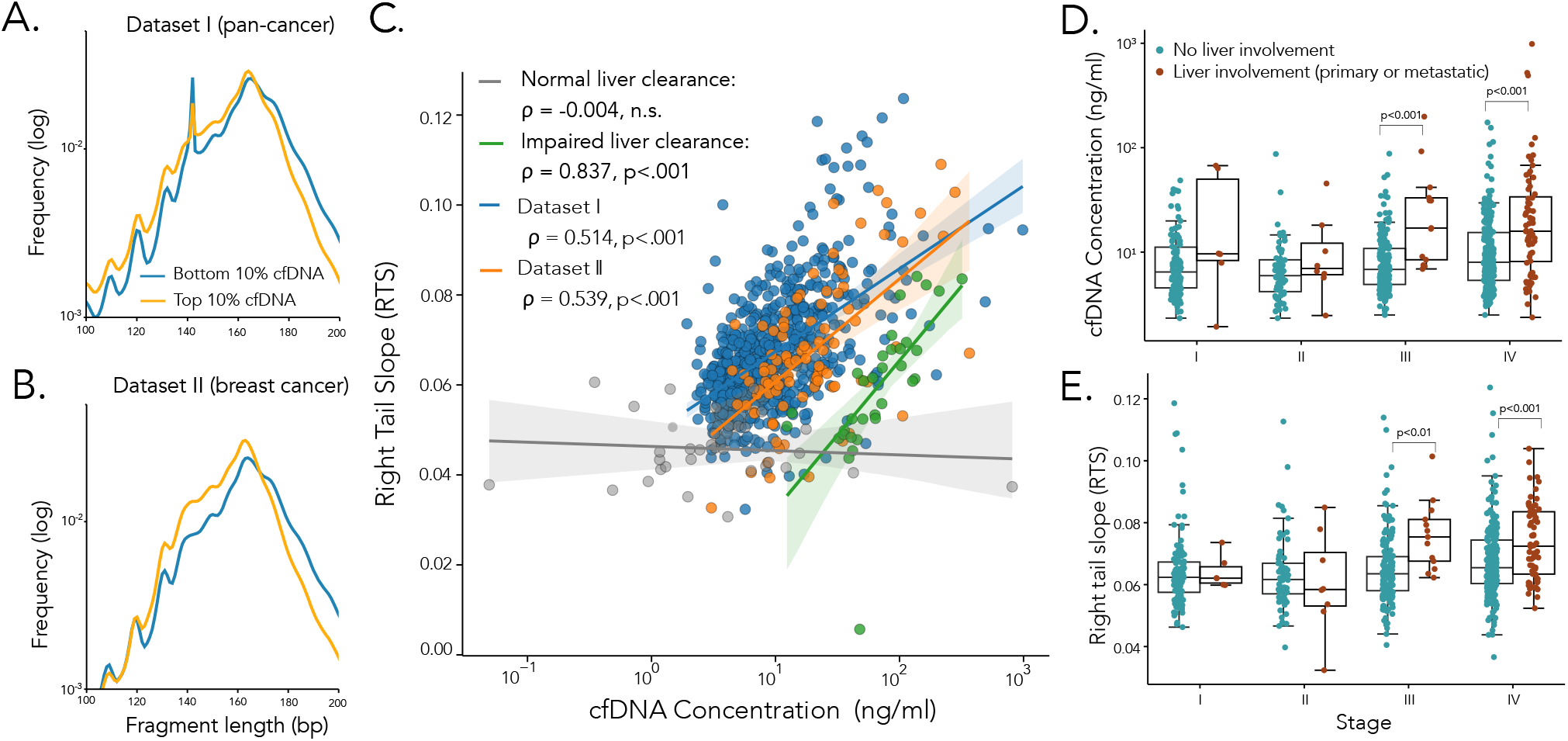
Comparing fragmentation in clinical samples to clearance experiments. **A-B:** Aggregate cfDNA distributions for human cancer patients in the top and bottom decile by cfDNA concentration for each dataset. **C** Association of cfDNA with RTS for Dataset I (blue), Dataset II (orange), post-nanoparticle-treated mice (green) and pre-nanoparticle-treated mice (gray). Bands represent the bootstrapped 95% CI. **D-E:** CfDNA concentration (**D**) and RTS (**E**) from Dataset I stratified by stage and liver involvement. Liver involvement includes samples from patients with primary liver cancer, primary hepatobiliary cancer, or liver metastases. P-values shown are for pairwise Mann-Whitney U tests corrected for stage-level comparisons via Benjamin-Hochberg with a FDR of 5%.

### Drift in cfDNA fragment length mode is associated with clinical outcome and prognosis

Along with the changes to the RTS of the fragment length distribution with nanoparticle treatment and cfDNA concentration, we also observe a slight decrease in mode fragment length with concentration (**Figure 4A-B**), which is highly correlated with the RTS (**Figure 5A**). We also detect a downward shift among the nanoparticle-treated mice which we do not observe in the control group (**Figure 5B**) and a decrease in mode with cfDNA concentration for nanoparticle-treated mice but not antibody-treated mice (Spearman *ρ* =.47, *p* =.006, **Supplementary Figure S5A-B**). A weak association between mode and tumor fraction appears in Dataset I (*ρ* = −0.22, *p <*.001), but is not replicated in Dataset II (*ρ* = 0.04, *p* = 0.8, **Supplementary Figure S5C**), while patients with liver involvement also have a lower mode fragment length (**Supplementary Figure S5D**). Mechanistically, when we relax the assumption that histone-bound DNA is perfectly protected from slicing, allowing limited exposure rather than introducing a separate ctDNA population, the mode drifts downward as the fragmentation rate increases. Exposing two bases (rather than 24, as in the ctDNA model) recapitulates the 166–164 bp range observed in the data (**Figure 5A**).

**Figure 5:**
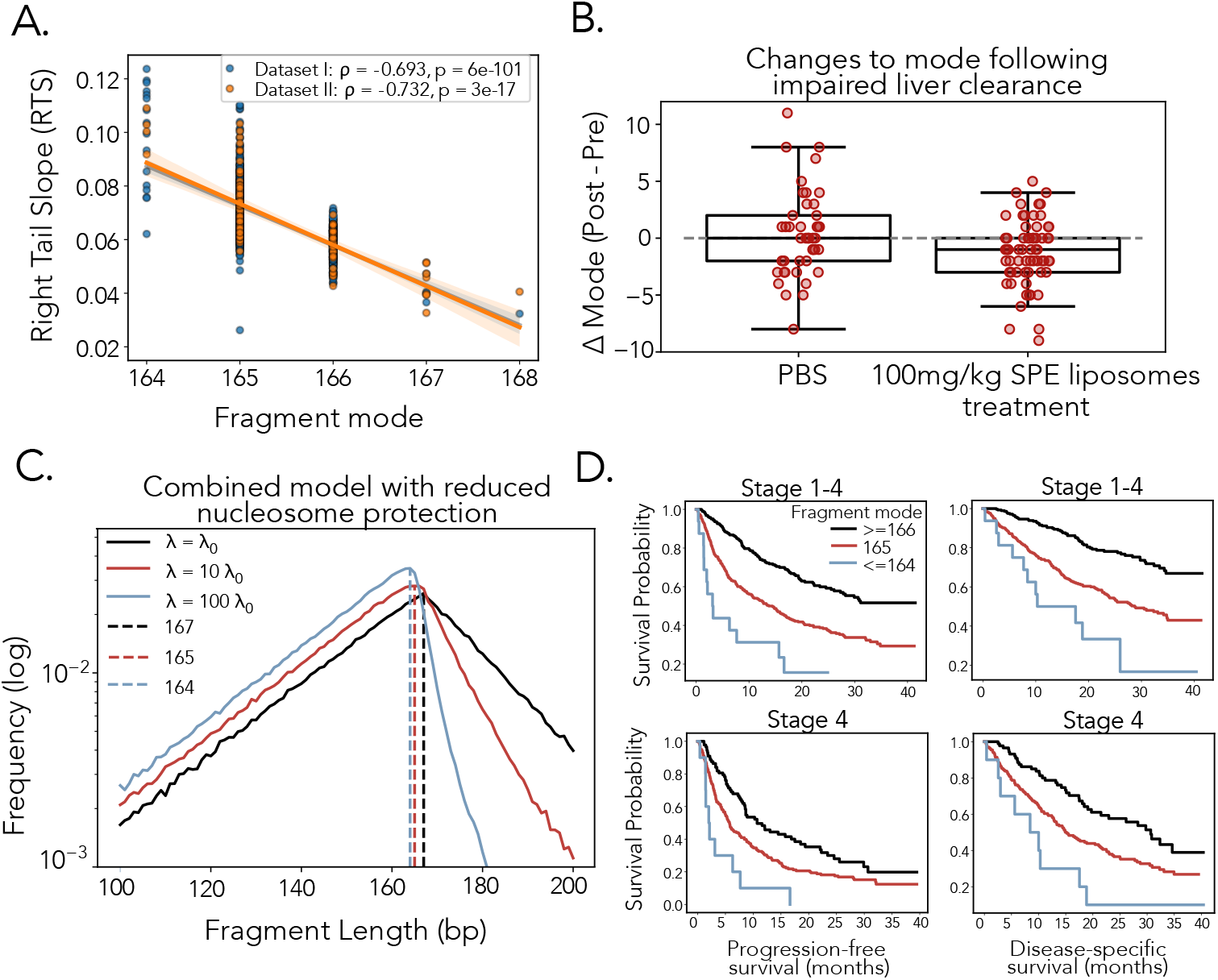
cfDNA fragment length mode and patient outcome. **A:** Correlation between *RTS* and mode fragment length for thw two patiet cohorts. **B:** Pairwise difference in mode fragment length for MA dataset mice, before and after treatment with PBS or SPE. **C:** The same simulation setup as **Figure 2A**, except 2 basepairs of the 167bp protected region are now exposed, causing the mode to drift left. **D:** Kaplan-Meier curves over progression-free and disease-specific survival for different mode fragment lengths for Stages I-IV and Stage IV only patients (in Dataset I). Results of multivariate Cox regression for each model are shown in **Table 1**.

Finally, we evaluated the association between fragment-based metrics and clinical outcomes in Dataset I using multivariable penalized Cox regression, for which progression-free and disease-specific survival data were available (**Table 1**). Neither RTS nor LTS was significantly associated with progression-free or disease-specific survival. However, stratification by modal fragment length revealed substantial differences in both outcomes, which remained significant after adjustment for cfDNA concentration, cancer stage, and MSAF across all four models (**Figure 5C**, **Table 1**).

**Table 1:**
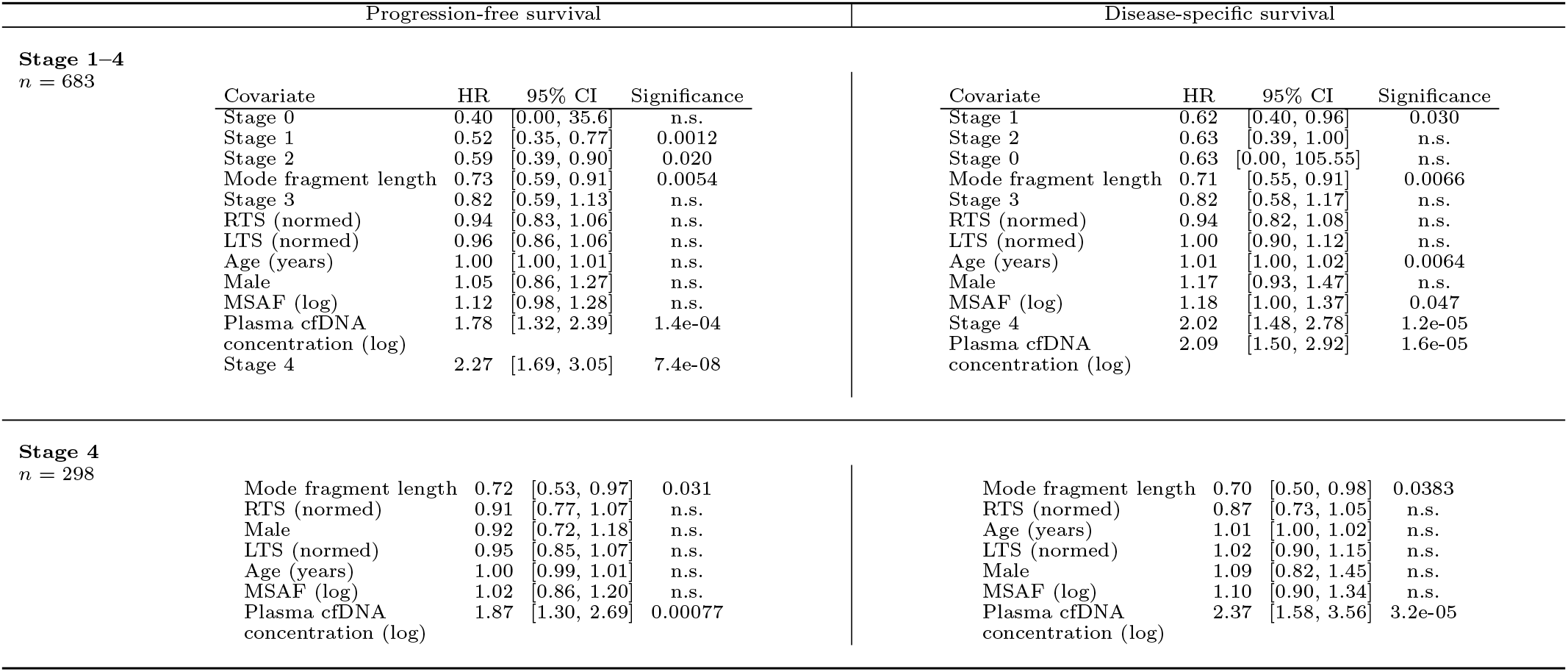
Cox model summaries by stage and endpoint.

| Progression-free survival |  |  |  |  | Disease-specific survival |  |  |  |
| --- | --- | --- | --- | --- | --- | --- | --- | --- |
| **Stage 1–4**<br>*n* = 683 |  |  |  |  |  |  |  |  |
| Covariate | HR | 95% CI | Significance |  | Covariate | HR | 95% CI | Significance |
| Stage 0 | 0.40 | [0.00, 35.6] | n.s. |  | Stage 1 | 0.62 | [0.40, 0.96] | 0.030 |
| Stage 1 | 0.52 | [0.35, 0.77] | 0.0012 |  | Stage 2 | 0.63 | [0.39, 1.00] | n.s. |
| Stage 2 | 0.59 | [0.39, 0.90] | 0.020 |  | Stage 0 | 0.63 | [0.00, 105.55] | n.s. |
| Mode fragment length | 0.73 | [0.59, 0.91] | 0.0054 |  | Mode fragment length | 0.71 | [0.55, 0.91] | 0.0066 |
| Stage 3 | 0.82 | [0.59, 1.13] | n.s. |  | Stage 3 | 0.82 | [0.58, 1.17] | n.s. |
| RTS (normed) | 0.94 | [0.83, 1.06] | n.s. |  | RTS (normed) | 0.94 | [0.82, 1.08] | n.s. |
| LTS (normed) | 0.96 | [0.86, 1.06] | n.s. |  | LTS (normed) | 1.00 | [0.90, 1.12] | n.s. |
| Age (years) | 1.00 | [1.00, 1.01] | n.s. |  | Age (years) | 1.01 | [1.00, 1.02] | 0.0064 |
| Male | 1.05 | [0.86, 1.27] | n.s. |  | Male | 1.17 | [0.93, 1.47] | n.s. |
| MSAF (log) | 1.12 | [0.98, 1.28] | n.s. |  | MSAF (log) | 1.18 | [1.00, 1.37] | 0.047 |
| Plasma cfDNA concentration (log) | 1.78 | [1.32, 2.39] | 1.4e-04 |  | Stage 4 | 2.02 | [1.48, 2.78] | 1.2e-05 |
| Stage 4 | 2.27 | [1.69, 3.05] | 7.4e-08 |  | Plasma cfDNA concentration (log) | 2.09 | [1.50, 2.92] | 1.6e-05 |
| **Stage 4**<br>*n* = 298 |  |  |  |  |  |  |  |  |
| Mode fragment length | 0.72 | [0.53, 0.97] | 0.031 |  | Mode fragment length | 0.70 | [0.50, 0.98] | 0.0383 |
| RTS (normed) | 0.91 | [0.77, 1.07] | n.s. |  | RTS (normed) | 0.87 | [0.73, 1.05] | n.s. |
| Male | 0.92 | [0.72, 1.18] | n.s. |  | Age (years) | 1.01 | [1.00, 1.02] | n.s. |
| LTS (normed) | 0.95 | [0.85, 1.07] | n.s. |  | LTS (normed) | 1.02 | [0.90, 1.15] | n.s. |
| Age (years) | 1.00 | [0.99, 1.01] | n.s. |  | Male | 1.09 | [0.82, 1.45] | n.s. |
| MSAF (log) | 1.02 | [0.86, 1.20] | n.s. |  | MSAF (log) | 1.10 | [0.90, 1.34] | n.s. |
| Plasma cfDNA concentration (log) | 1.87 | [1.30, 2.69] | 0.00077 |  | Plasma cfDNA concentration (log) | 2.37 | [1.58, 3.56] | 3.2e-05 |

## Discussion

Here, we show that a mechanistic model of saturating cfDNA clearance from the bloodstream reproduces key features observed across multiple liquid biopsy datasets, including previously published mouse models that directly perturb clearance pathways and newly analyzed plasma samples from patients with primary and metastatic cancer. In this framework, hepatic clearance of cfDNA saturates rather than increasing linearly with concentration, leading to prolonged circulation times that provide a parsimonious explanation for both elevated cfDNA levels and enhanced fragmentation reported in prior studies. By explicitly modeling DNA shedding into the circulation, clearance saturation, and fragmentation, this approach enables the inference of independent and interpretable features of these processes from observed data. Importantly, these features provide prognostic information beyond cfDNA concentration alone, indicating that fragment length distributions capture biologically distinct aspects of cfDNA dynamics implicated in disease progression.

While minimal mechanistic models are powerful for isolating core biological mechanisms and providing intuitive insight into system behavior, they necessarily abstract away additional layers of biological complexity. One alternative explanation for the observed fragmentation patterns is that elevated cfDNA directly induces nuclease activity, leading to increased fragmentation independent of clearance saturation or circulation time, and preferentially targeting exposed DNA [34]. However, this hypothesis predicts that fragmentation features should scale directly with cfDNA concentration, including in pre-treatment, normal liver clearance mouse samples, which we do not observe (**Figure 4C**). Moreover, whether increased fragmentation arises from a higher fragmentation rate or from prolonged exposure does not alter the central relationship predicted by our model between reduced hepatic clearance and fragment length. Together, these considerations support clearance saturation and extended circulation time as the most parsimonious explanation consistent with the available data.

Although increased circulation time is likely a central driver of the observed fragmentation patterns, further work is needed to explain why fragment length distributions in both mice with impaired hepatic clearance and patients with high cfDNA levels are better explained by the slicing model of nucleosome fragmentation, rather than by a concurrent increase in both forms of fragmentation, as might be expected from a global reduction in clearance. One possible explanation is that fragmentation rates differ over time across cutting mechanisms: circulating nucleases may cleave nucleosome-bound DNA only a limited number of times before destabilizing nucleosome association, or nucleosomes may progressively dissociate from DNA during circulation. Under these conditions, impaired clearance would prolong fragment persistence in the bloodstream, increasing opportunities for linker slicing, while the resulting shorter nucleosome-protected fragments may be underrepresented in standard double-stranded DNA libraries. Consequently, prolonged circulation time, without changes in nucleosome stability, would preferentially enrich fragmentation signatures on exposed linker DNA, consistent with reports that library preparation methods optimized for single-stranded DNA detect substantially more short cfDNA fragments [30, 35].

It is notable that aST3, which binds to cfDNA, predominantly affects the left tail of the fragment length distribution rather than the right. One possible explanation is that interactions with aST3, previously shown to stabilize nucleosomes, allow cfDNA fragments to remain nucleosome-bound long enough to undergo additional fragmentation that would otherwise render them undetectable. Under this hypothesis, inhibition of enzymatic clearance permits fragmentation to accumulate while cfDNA remains protected by nucleosomes. As expected under prolonged circulation time, the ratio of LTS before and after treatment predicts the observed fold-chage of cfDNA (**Supplementary Figure S6**). Although our results show that increased trimming captures the net effect of antibody priming, as we might expect if it acts in part by prolonging protection of nucleosome-bound DNA, a complete mechanistic description of this process lies beyond the scope of this study. Nevertheless, antibody priming produces large and predictable shifts in the abundance of short cfDNA fragments, suggesting that therapeutic strategies aimed at increasing cfDNA yield through this approach may complicate fragmentomics-based prediction. In contrast, interference with Kupffer cell–mediated clearance has a more limited effect on fragments below 160 bp, although biologically informative signals may still be obscured by a reduction in the modal fragment length. Given the clinical relevance of inflammation across many disease contexts and its demonstrated impact on cfDNA fragmentation patterns, caution is warranted when interpreting fragment-based biomarkers [36].

We provide a mechanistic explanation for the decrease in modal fragment length that is independent of circulating tumor DNA (ctDNA) content and show that this signature is associated with cancer survival; however, the biological basis of this association remains uncertain. Tumor growth may simultaneously impair clearance capacity and increase cfDNA shedding, but if ctDNA were the primary driver of increased shedding, one would expect a strong association between fragmentation features and tumor fraction, which we do not observe. This finding suggests that non-tumor cells contribute substantially to elevated cfDNA levels. Such increases may still arise indirectly from tumor growth, for example through inflammation or damage to healthy tissue [37, 36]. Consistent with this possibility, tissue-specific histone modifications have been shown to increase with cfDNA concentration [38]. Elevated levels of cfDNA bearing modified histones may also increase exposed linker DNA available for fragmentation and decrease the mode fragment length, consistent with our adapted model (**Figure 5**). Alternatively, high-risk patients may be more likely to present with comorbidities—such as hypertension, obstructive sleep apnea, or chronic inflammation—that independently elevate cfDNA shedding and are themselves associated with poor outcomes [39, 40, 41, 42].

This study has several limitations that define the scope of our conclusions. First, our model does not explicitly incorporate tumor-specific fragmentation dynamics, which may contribute additional differences between tumor-derived and non–tumorderived cfDNA, particularly at high concentrations. Second, we do not directly compare patient or mouse data to a healthy cohort. However, tumor fractions in the mouse experiments were uniformly low and decreased slightly following each treatment in the original analysis, so that ctDNA was unlikely to confound observed effects on fragment length. Given the magnitude of the observed shifts in fragment length distributions, similar effects would be expected in non–tumor-bearing mice. Notably, a recent study of cfDNA variation in healthy individuals reported that high cfDNA burden was associated with reduced nuclease activity and increased fragment lengths, particularly through enrichment of the second nucleosome wrap [43]. Rather than contradicting our findings, this pattern suggests that variation in healthy populations may reflect reduced enzymatic clearance instead of impaired hepatic clearance, consistent with a lower slicing rate. That this behavior contrasts with the effects of antibody priming, intended to inhibit enzymatic clearance, further supports the view that antibody priming does not eliminate but rather alters nuclease activity. Together, these results indicate that healthy and diseased states may involve distinct modes of clearance impairment with different consequences for cfDNA fragmentation. Future work should leverage mechanistic fragmentation models to distinguish these processes and to better understand the origins of elevated cfDNA across disease contexts beyond cancer.

Overall, we show that saturation of hepatic clearance contributes to elevated cfDNA concentrations in cancer and promotes fragmentation of exposed linker DNA. This process is strongly associated with a reduction in modal fragment length, a feature that carries prognostic information independent of cfDNA concentration. Although tumor-derived cfDNA contributes a distinct signal, it does not explain these fragmentation patterns or their prognostic value. More generally, our findings demonstrate how mechanistic modeling can extract biologically meaningful and clinically relevant insights from data that may appear noisy or limited, where a purely data-driven statistical or machine learning approach will fail. Mechanistic frameworks of this kind offer a principled foundation for interpreting cfDNA fragmentomics and may improve its reliability for clinical inference in cancer and other high–cfDNA-burden diseases.

## Code and Data availability

All data needed to evaluate the conclusions in the paper are present in the paper and the Supplementary Materials or are included with the source code described below. Additional data related to this paper may be requested from the authors. Custom scripts were used for the simulations and data analyses and all code is available on Github at https://github.com/trachman1/saturatingClearance2026. All packages used for analysis and visualization are open-source.

## Acknowledgments

This research was done using resources provided by the Open Science Grid, which is supported by the National Science Foundation award 1148698, and the U.S. Department of Energy’s Office of Science.

## Materials and Methods

### Model calibration

#### Calibrating *s, k, n*_0_, *l*_0_

The ratio of *s* to *k* determines the number of fragments at equilibrium, but do not independently influence the fragment distribution. To create a sufficiently large fragment sample, we set *s* = 10^8^ and *k* = 200. By default, the protected region *n*_0_ is set to 167bp and *l*_0_ is set to 0.

#### Calibrating *v, λ, β*

Based on our analytical model, the identifiable parameters are the ratio of *k* to both forms of fragmentation. The value 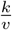 is equivalent to LTS. When *k >> λ*, the RTS is approximately the exponential rate *β* that sets the initial linker length distribution. Otherwise, it is monotonically related to the ratio 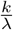. To calibrate the model to fragment distributions we first use control samples (PBS and Isotype treatments) in the MA dataset, which we assume to have low fragmentation rates relative to clearance. We therefore set the length parameter *β* to the median RTS in this group and the trim rate *v* to the ratio of the (arbitrary) clearance rate *k* to the median LTS. Because the median LTS and RTS are both approximately 0.043, we set *β* = 0.043 and *v* = 200*/*0.043 = 4651. For the ctDNA model (**Figure 3**), we calibrate the combined model to clinical samples by setting the baseline LTS to minimize the effect of ctDNA, setting 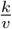 to the median LTS among samples with an MSAF below 0.1% in Dataset I, which is 0.056. To avoid the effects of concentration on RTS in calibrating *β*, we set it to the median RTS among the bottom decile by cfDNA concentration in Dataset I, which is 0.059.

#### Mode estimation

We estimate the mode as the maximum frequency length. To consistently measure the mode of the peak near 167bp, rather than a secondary peak, we consider lengths exceeding 160bp.

### Combined model

To study the effect of both slicing and trimming on cfDNA lengths, we combined both processes in one simulation. We implement a continuous-time stochastic simulation of cfDNA fragmentation and clearance using a Gillespie algorithm [44]. Each cfDNA fragment is modeled as a single nucleosome flanked by two linker regions and is represented by the state

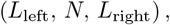

where *L*_left_ ≥ 0 and *L*_right_ ≥ 0 denote the lengths (in base pairs) of the left and right linker DNA, respectively, and *N* ≥ 0 denotes the remaining nucleosomal DNA length. Fragments enter the system as a Poisson process with rate *s*. Upon entry, the nucleosome length is initialized to *N* = *n*_0_, and the total linker length is initialized as

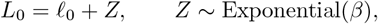

where *l*_0_ is a baseline linker length and *β* is the exponential rate parameter. The total linker length *L*_0_ is split uniformly at random between the left and right linkers. Fragment evolution proceeds via three additional event types: linker slicing, end trimming, and clearance. Linker slicing occurs at a per-base-pair rate *λ*_linker_, yielding a total fragmentation propensity

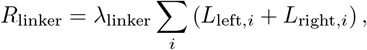

where the sum is taken over all active fragments. Conditional on a linker fragmentation event, a fragment is selected with probability proportional to its total linker length, and a cut position is drawn uniformly along that fragment’s linker DNA. The outer fragment produced by the cut is discarded, and only the nucleosome-containing fragment is retained, resulting in a reduction of the linker length. Trimming occurs at rate *v* per fragment. For each trimming event, a fragment is selected uniformly at random, and the left or right end is chosen with probability 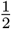. A single basepair is removed from that end, first consuming linker DNA and then nucleosomal DNA if necessary. If trimming completely removes the nucleosome (*N* ≤ 0), the fragment is removed from the system. Fragments are cleared independently at Poisson-rate *k* per fragment. To efficiently sample fragments in proportion to their linker lengths during fragmentation events, we maintain a binary-indexed tree over per-fragment weights

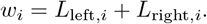

This data structure enables sampling and updating to the fragment pool in 𝒪(log *n*) time, where *n* is the number of active fragments. The total linker length Σ_*i*_ *w*_*i*_ is tracked explicitly to allow constant-time computation of slicing propensity. We simulate both continuous and discrete base-pair-resolution cutting and observe minimal differences in the stationary distributions (**Supplementary Figure S7**).

#### Exposed linker model

To study the effect of leaving some of the nuclosome wrap open to fragmentation, we allow *l*_0_ > 0. To keep the total initial fragment length consistent, we update *n*_0_ ← 167 − *l*_0_. In **Figure 3** of the main text, we set *n*_0_ = 143 bp, *l*_0_ = 24 bp to mimic ctDNA. In **Figure 5**, we use the same model with *n*_0_ = 165 bp, *l*_0_ = 2 bp.

#### ctDNA Mixture Model

We define the mixture of tumor and non-tumor length distributions as a standard mixture model. Given tumor and normal cfDNA length distributions *f* (*x*) and *g*(*x*), for a tumor fraction *p* ∈ [0, 1], the mixture is

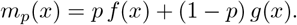

### Martin-Alonso Dataset

All SAM files and metadata from Martin-Alonso et al. were downloaded from SRA (ID: PRJNA1037081). CfDNA concentrations were taken from supplementary data provided in [23]. The mice in Supplemental Section S3 were used for concentration data for the nanoparticle treatment. Additional read data available in SRA from mice treated with PBS or 100mg/kg SPE liposomes were included for analyses that did not require concentrations including both retro-orbital and terminal bleed samples. The same analyses were run on only retro-orbital samples with similar results (**Supplemental Figure S8**). The samples in Section S5 and corresponding reads were used to study the effect of aST3 antibody treatment. We detail the samples used for each analysis in **Supplementary Table S9**.

### Patient Cohorts

Dataset I was compiled from individuals enrolled in the Moonshot study at Allegheny Health Network. Clinical information was available for 874 patients and plasma sequencing from a single blood draw for 716 patients. The intersection of these was a total of 696 patients. A breakdown of clinical characteristics is given in **Supplementary Tables S1-S2**.

Dataset II consisted of 112 samples taken from 31 individuals with metastatic breast cancer. Concentration, tumor fraction, and fragmentomic data were available for 95 total samples from the 31 individuals, which were used in all analyses. All patients had HR+HER2-metastatic invasive lobular breast cancer (ILC) and were included in the University of Pittsburgh Breast Disease Research Repository (HCC 04-162). A breakdown of cfDNA and tumor fraction by patient is given in **Supplementary Table S3**.

### Sample preparation and sequencing

#### Dataset I

Plasma samples from Dataset I were purified and sequenced using the TruSight Oncology 500 (TSO500) assay following the previously published Materials and Methods in [32] and **Supplementary Methods 1.2**.

#### Dataset II

Plasma samples from Dataset II were purified and then sequenced using ultra-low-pass whole-genome sequencing (ULP-WGS). For full information on consent and sample collection, plasma isolation, cfDNA purification, sequencing, and analysis, see **Supplementary Methods 1.3**.

### Bioinformatics

#### Fragmentomics

BAM files were filtered for mapping quality and properly paired reads using samtools view. cfDNA fragment lengths were computed from each BAM file as the insert size of the mapped read. Fragment length densities were normalized to the 100-200bp range to select for mononucleosome-length fragments. A sample was dropped if after post-processing it did not have at least one fragment per bin in the 100-200bp range.

#### Tumor fraction estimation

For Dataset I, the tumor fraction was estimated from the Maximum Somatic Allele Frequency (MSAF) based on the DRAGEN TSO500 ctDNA Analysis pipeline [45]. Samples in Dataset II underwent whole-genome sequencing and so IchorCNA was used to assess tumor fraction [46]. For more accurate comparison of the two quantities, for Datset I the tumor fraction was considered to be 2*MSAF, assuming the MSAF is the frequency of a haploid clonal variant.

### Statistical analysis

All statistical analyses were performed in Python. Paired comparisons were assessed using the Wilcoxon signed-rank test (scipy.stats.wilcoxon) and non-paired comparisons were performed using a Mann-Whitney U test with post-hoc corrections using the Benjamin-Hochberg procedure with a false discovery rate of 5% (pingouin.pairwise tests). Associations were evaluated using Spearman correlation coefficients (scipy.stats.spearmanr) by default; Pearson correlation coefficients were used when testing for linear relationships. Cox proportional hazards models and Kaplan–Meier survival curves were fit using the lifelines package. Between-group differences in Kaplan–Meier curves were assessed using the log-rank test. Cox models were fit with an *L*_2_ penalty of 0.1 to mitigate multicollinearity. A two-sided p-value threshold of 0.05 was used to determine statistical significance.

