## Supplementary Material for "Saturating hepatic clearance drives elevated cfDNA and fragment shortening in cancer"

**2 Supplementary Figures** **13**
**3 Supplementary Tables** **20**

### 1 Supplementary Methods

#### 1.1 Mathematical model of cfDNA fragmentation under saturating clearance

##### 1.1.1 cfDNA yield dynamics

Let $C(t)$ denote the circulating cfDNA concentration, $s$ the shedding rate into circulation, and $k$ the clearance rate from the blood. Under a linear clearance assumption, cfDNA dynamics are governed by

$$\frac{dC}{dt} = s - kC, \tag{1}$$

such that, at steady state, the equilibrium cfDNA concentration is proportional to the ratio of shedding into the blood to clearance from the blood,

$$C^* = \frac{s}{k}. \tag{2}$$

To model clearance saturation, we instead assume cfDNA removal follows Michaelis–Menten kinetics,

$$\frac{dC}{dt} = s - \frac{V_{\max}C}{K_m + C}, \tag{3}$$

where $V_{max}$ denotes the maximal clearance capacity and $K_m$ the half-saturation constant. At steady state, the cfDNA concentration is

$$C^* = \frac{K_m S}{V_{\max} - s}. \tag{4}$$

In this regime, the effective per-molecule clearance rate is $\frac{V_{max}-s}{K_m}$, which decreases toward zero as the shedding rate and tends toward zero as $s$ approaches $V_{max}$. Because inflow and outflow are equal at steady state, the mean residence time of cfDNA in circulation is given by Little's theorem as

$$\frac{C^*}{s} = \frac{K_m}{V_{\max} - s}, \tag{5}$$

which diverges as clearance capacity is approached [1].

To model tumor growth, we decompose the total shedding rate as $s = s_N N(t) + s_T T(t)$, where $N(t)$ and $T(t)$ denote the numbers of non-tumor and tumor cells, respectively, and $s_N$ and $s_T$ are their corresponding per-cell shedding rates.

While in the example shown in **Figure 1B** we assume $N(t)$ is constant, non-tumor cfDNA may primarily drive cfDNA burden in the case of inflammation. In our illustration, $s_N = s_T = 10^-5$ molecules $*$ cells $^{-1}*$ time$^{-1}$ , $k = 1$ time$^{-1}$, $N = 10^9$ cells, and $T$ ranges from 10 to $10^{10}$ cells. To show behavior near $V_{max}$, we set it to $1.01 * \max\{s\}$, and to keep effective clearance equivalent to $k$ for small $C$, we set $K_m = V_{max}$ molecules.

##### 1.1.2 cfDNA fragmentation dynamics

We model cfDNA entering circulation as mononucleosomal fragments. This choice reflects the use of double-stranded DNA library preparations optimized for mononucleosomal fragment detection, which represent the most abundant and reliably detectable fragment size range in plasma.

We define the initial mononucleosome fragment size as having minimum length $n_0$ and additional base pairs drawn from an exponential distribution, so that

$$x_0 = n_0 + Z,\, Z \sim Expo(\beta).$$

By default, we set $n_0 = 167$ bp, corresponding to a 143 bp nucleosome core particle with an additional 24 bp associated with the linker histone. We choose an exponential distribution for the linker length based on the approximately linear behavior of the right tail of the mononucleosome peak in log-frequency space, and to maintain consistency with prior modeling work [2].

We allow two fragmentation mechanisms: slicing and trimming. Slicing applies only to exposed linker segments while trimming applies to the entire fragment.

##### 1.1.3 Slicing model

Linker fragments are characterized only by their total length $x > 0$. New fragments enter as a Poisson process of rate $s$, with i.i.d. entry lengths drawn from a density $p_0(x)$. Each fragment of length $x$ undergoes fragmentation at length-dependent rate $\lambda x$ (per-bp cut rate $\lambda$), and conditional on fragmentation, a cut position is chosen uniformly along the fragment. Only the left daughter is retained, which is assumed to be attached to the nucleosome core particle. The right daughter fragment is dropped. Fragments are cleared at constant per-fragment rate $k$.

Let $f(x)$ denote the stationary density of fragments of length $x$. Define the stationary tail mass

$$S(x) \;=\; \int_x^\infty f(u)\,du, \qquad \text{so that} \qquad f(x) = -S'(x).$$

**1.1.3.1 Stationary balance equation.** At equilibrium, for each $x > 0$ the net flux into length $x$ is zero:

$$0 \;=\; s\,p_0(x) \;+\; \underbrace{\lambda\int_x^\infty f(u)\,du}_{\text{gain from fragmentation of longer fragments}} \;-\; \underbrace{(\lambda x + k)\,f(x)}_{\text{loss from fragmentation and clearance}}\;.$$

The gain term is $\lambda S(x)$ because a fragment of length $u$ fragments at rate $\lambda u$ and produces a left-daughter of length in $[x, x+dx]$ with probability $dx/u$, yielding $\lambda f(u)dx$ after cancellation and integration over $u \geq x$.

Substituting $f(x) = -S'(x)$ gives a first-order linear ODE for $S$:

$$\lambda S(x) \;=\; (\lambda x + k)(-S'(x)) \;-\; s\,p_0(x),$$

or equivalently

$$(\lambda x + k)S'(x) + \lambda S(x) = -s\,p_0(x).$$

**1.1.3.2 Closed-form solution for general entry distribution.** Divide by $(\lambda x + k)$:

$$S'(x) + \frac{\lambda}{\lambda x + k}S(x) = -\frac{s\,p_0(x)}{\lambda x + k}.$$

The integrating factor is

$$\mu(x) = \exp\!\left(\int \frac{\lambda}{\lambda x + k}\,dx\right) = \exp(\ln(\lambda x + k)) = \lambda x + k.$$

Thus,

$$\frac{d}{dx}\Big[(\lambda x + k)S(x)\Big] = -s\,p_0(x).$$

Using the boundary condition $S(\infty) = 0$ (finite stationary mass), integrate from $x$ to $\infty$:

$$(\lambda x + k)S(x) = s \int_x^\infty p_0(u)\, du = s\, \overline{F}_0(x),$$

where $\overline{F}_0(x)$ is the survival function of the entry distribution. Therefore,

$$S(x) = \frac{s\, \overline{F}_0(x)}{\lambda x + k}.$$

Making use of the fact $f(x) = -S'(x)$ and differentiating $S(x)$ yields a general expression:

$$f(x) = s \left[ \frac{p_0(x)}{\lambda x + k} + \frac{\lambda\, \overline{F}_0(x)}{(\lambda x + k)^2} \right],$$

which we can see becomes the initial linker distribution $p_0$ as $\lambda \to 0$. The expected total number of fragments at stationarity is $\int_0^\infty f(x)\, dx = S(0)$, giving

$$\int_0^\infty f(x)\, dx = S(0) = \frac{s\, \overline{F}_0(0)}{k} = \frac{s}{k},$$

as expected. The general form of the normalized fragment profile $\frac{k}{s} f(x)$ for $\lambda > 0$ is then

$$\frac{k}{\lambda} \left[ \frac{p_0(x)}{x + \frac{k}{\lambda}} + \frac{\overline{F}_0(x)}{(x + \frac{k}{\lambda})^2} \right].$$

**1.1.3.3 Exponential entry.** Let the entry length be exponential with rate $\beta > 0$:

$$p_0(x) = \beta e^{-\beta x}, \qquad \overline{F}_0(x) = e^{-\beta x}.$$

Then the stationary tail mass is

$$S(x) = \frac{s\, e^{-\beta x}}{\lambda x + k},$$

and the normalized stationary density for $\lambda > 0$ is

$$\frac{k}{\lambda} e^{-\beta x} \left[ \frac{\beta}{x + \frac{k}{\lambda}} + \frac{1}{(x + \frac{k}{\lambda})^2} \right].$$

##### 1.1.4 Trimming model

In circulating plasma, nucleases preferentially cut histone-bound DNA at outward-facing minor grooves, producing characteristic small peaks in fragment length distributions at 10.4 bp intervals. Rather than model this oscillation, we focus on the net increase of dsDNA fragments in this range until the length of the mononucleosome wrap. We approximate this process using a constant deterministic erosion of fragment ends at rate $v$ ( $\mathrm{bp\,time}^{-1}$).

Let $x_0$ denote the initial fragment length at entry. After time $T$, the fragment length is

$$x(T) = \max\{0, x_0 - vT\}.$$

Based on an assumption that short nucleosome-bound fragments are cleared at identical rates independent of the exact fragment length, clearance is modeled as a Poisson process with constant rate $k$, so that

$$T \sim \text{Exponential}(k).$$

Rescale the waiting time in terms of basepairs consumed

$$U = vT,$$

so that $U \sim \text{Exponential}(c)$ with rate

$$c = \frac{k}{v}.$$

Then the fragment length conditional on initial total length $x_0$ is

$$f(x|x_0) = U(x_0 - x).$$

We assume that initial fragments consist of a protected nucleosomal region of fixed length $n_0$, plus an exposed region whose length is exponentially distributed. Thus,

$$x_0 = n_0 + Z, \qquad Z \sim \text{Exponential}(\beta),$$

with probability density

$$f_{x_0}(x) = \beta\, e^{-\beta(x-n_0)}\, \mathbf{1}_{\{x \geq n_0\}}.$$

We now solve $f(x)$ for $x > 0$. Conditioning on $x_0$ and $U$,

$$f(x) = \int f(x|x_0) f_{x_0}(x_0) = \int_{-\infty}^{\infty} f_{x_0}(x_0)\, U(x - x_0)\, dx_0.$$

Because the initial fragment cannot be less than $n_0$, we evaluate the integral from $\max(x, n_0)$. Substituting the densities yields

$$f(x) = \int_{\max(x,n_0)}^{\infty} \left(\beta\, e^{-\beta(x_0-n_0)}\right)\left(c\, e^{-c(x_0-x)}\right) dx_0.$$

Factoring out terms independent of $x_0$, we can write

$$f(x) = \beta c\, e^{\beta n_0 + cx} \int_{\max(x,n_0)}^{\infty} e^{-(\beta+c)x_0}\, dx_0 = \frac{\beta c}{\beta + c}\, e^{\beta n_0 + cx}\, e^{-(\beta+c)\max(x,n_0)}.$$

**1.1.4.1 Piecewise form.** Splitting by whether erosion has reached the nucleosome boundary yields the following expression

$$f_{trim}(x) = \begin{cases} \dfrac{\beta c}{\beta + c}\, e^{\,c\,(x-n_0)}, & 0 < x < n_0, \\[2ex] \dfrac{\beta c}{\beta + c}\, e^{-\beta\,(x-n_0)}, & x \geq n_0. \end{cases}$$

#### 1.2 Dataset I (AHN Moonshot) Supplementary Materials and Methods

**Note:** This supplement is adapted directly from [3] but contains information specific to the current publication.

##### 1.2.1 Consent and Sample Collection.

This study included male and female patients (age 18 to 100) diagnosed with cancer of any origin who came to AHN for clinical care. Participants signed a HIPAA Authorization Statement of informed consent (IRB# 2020-258: Oncology Sample Biobank and Data Repository) to contribute de-identified data to a database linking their tumor and blood sequencing data with clinicopathological tumor information. The protocol included access to cancer patients having procedures at one of 21 AHNCI treatment sites including provision of blood samples obtained during routine, clinical lab draws. Two-thirds of the patients selected for this study had samples acquired at or near the time of initial diagnosis and prior to any interventional therapy. Whole blood samples were collected in three, 10 mL Streck, Cell-Free DNA BCT tubes (Streck, LaVista, NE) according to a standard operating procedure specifying gentle inversion of each tube ten times after drawing, wrapping each tube in bubble wrap, and maintenance of samples at room temperature (18oC to 25oC). Streck tubes were transferred daily by medical courier to the AHN Genomics Facility for processing. Samples subjected to undue agitation including pneumatic tube system transport were not included in the study.

##### 1.2.2 Isolation of Plasma from Streck Tube Samples.

We developed a 3-step sequential centrifugation protocol to separate plasma devoid of cells and debris from patient blood samples. Whole blood in Streck tubes was gently mixed and placed in a chilled, swinging bucket rotor for centrifugation at

1600 rcf (Eppendorf 5810R, S-4-104 rotor, 10 min, 4oC, no brake, Eppendorf, Hamburg, GER). All blood products remained on ice or cold blocks (15 ml x 12, Electron Microscopy Sciences, Hatfield, PA) from the first spin throughout the plasma and buffy coat separation. The plasma layer obtained after centrifugation was transferred to a 5mL screw cap, conical tube (VWR, Inc., Radnor, PA) without disturbing the buffy coat using a 10 mL pipet tip (Rainin Pipet-Lite LTS L-10 mL, Mettler-Toledo, Greifensee, CH). The plasma was centrifuged in a fixed-angle rotor at 10,000 rcf (Eppendorf 5425R, FA 10 x 5 rotor, 10 min, 4oC, soft brake). The plasma layer was then transferred to a 5 mL screw-cap tube using a 10 mL serological pipet (cat. #14955234, Drummond pipet-aid XL, ThermoFisher,) without disturbing the cell pellet and a third spin was performed identically to the previous spin. After the third spin, the plasma was transferred to a 7.6 mL FluidX Tricode tube (Azenta, Burlington, MA) for immediate storage (-80oC) in the Biospecimen repository. The buffy coat layer separated in the initial Streck tube was transferred to a 1.9 mL FluidX Tricode tube (Azenta) using a large-bore, 1000 µL pipet tip (RT LTS 1000 µL, Mettler-Toledo) for long-term storage (-80oC).

##### 1.2.3 Purification of Cell Free DNA from Plasma.

Frozen plasma was thawed at room temperature (60 min) and the volume adjusted as needed (1X PBS, pH 7.4; cat. #10010-023, ThermoFisher) for purification of cell-free DNA (cfDNA) using the Apostle MiniMax High Efficiency cfDNA Isolation Kit (cat. #A17622-250, Beckman, Indianapolis, IN). The only protocol modification was the primary Proteinase K digestion incubation that we optimized to 1 hour at 60°C. Cell-free DNA was eluted using magnetic beads in 40-50 µL of elution buffer per sample and 2 µL was assayed for concentration by fluorometry (Qubit Flex, ThermoFisher) using the Qubit dsDNA High Sensitivity kit (cat. #Q32854, ThermoFisher). An aliquot was diluted to 200-600 pg/µL and 2 µL were analyzed on a 5200 Fragment Analyzer (Agilent, Santa Clara, CA) using the HS Large Fragment Kit (DNF-464, Agilent). The analysis software (ProSize Revision 5.0.1.6, Agilent) delineated the DNA fragment distribution from 75 bp-300 bp and 75 bp-1200 bp containing cell free DNA (cfDNA) of sizes associated with nucleosome and linker histones and the 1300 bp-150000 bp domain where "background" germline DNA from lysed cells was detected when present. The percentages of those regions were multiplied by Qubit Flex quantitation values to obtain the concentration of these variable DNA components for each sample.

##### 1.2.4 Sequencing Library Preparation of Cell Free DNA Samples.

Sequencing was performed on the initial blood draw obtained from 716 cancer patients for whom comprehensive clinical information including outcomes was available for 697 subjects. Libraries were prepared for sequencing cfDNA according to the Illumina TruSight Oncology 500 ctDNA Reference Guide (Doc. #1000000092559 v00 Feb. 2020, Illumina). Briefly, a minimum of 25-30 ng cfDNA contained within the 75bp-300 bp DNA domain was required without fragmentation. Library preparation included end repair, A-tailing, UMI adapter ligation and index PCR yielding a concentration of approximately 80-170 ng/µl. Two rounds of target enrichment were performed followed by PCR to obtain fragments in the 325-375 bp range at a concentration of 20 ng/µL. Twenty-four sample libraries were manually normalized (0.65 nM) and paired-end sequencing (2 X 150 bp) was performed on the NovaSeq 6000 according to the NovaSeq 6000 Sequencing System Guide (Doc. #1000000019358 v17 Sept. 2022, Illumina).

###### 1.2.5 Data Analysis Pipeline.

Cell free DNA libraries sequenced on the NovaSeq6000 (Illumina) were transferred to and processed in Illumina Connected Analytics (ICA) using the DRAGEN TruSight Oncology 500 ctDNA Analysis software v1.2 (Doc. 200015532 v00, Aug. 2022). Analysis was performed using the DRAGEN TSO500 ctDNA RUO v1.2 pipeline. The processing paradigm converted BCL files to FASTQ files using 1) the BCL Convert algorithm to demultiplex the raw data, 2) a UMI processing step to perform read collapsing and identify low frequency variants, 3) mapping and alignment to the hg19 reference genome using the DRAGEN aligner followed by 4) stretched realignment and paired read-stitching using GEMINI. All cell free DNA samples were processed identically using this analysis pipeline.

##### 1.3 Dataset II (breast cancer patients) Supplementary Materials and Methods

###### 1.3.1 Consent and sample collection

Participants provided written informed consent for inclusion in the University of Pittsburgh Breast Disease Research Repository (HCC 04-162), permitting collection, long-term storage, and research use of biospecimens and associated clinical data. Specimens were obtained during routine clinical care or at concurrent time points, with no additional surgical procedures performed. For this study, all biospecimens were standardized to peripheral blood collection and drawn into EDTA or Streck Cell-Free DNA BCT® tubes, with 10mL of blood in two separate tubes. Samples were coded, de-identified for research use, and stored under institutional oversight for future molecular analyses.

###### 1.3.2 Plasma isolation

This study was conducted under an Institutional Review Board–approved protocol (HCC 04-162 / IRB#21020005). Peripheral blood was collected during routine clinical visits into EDTA or Streck Cell-Free DNA BCT® blood collection tubes. Samples were processed following Pitt Biospecimen Core standard operating procedures for double-spun plasma preparation. Briefly, blood was maintained at room temperature and processed within 96 hours of collection. Whole blood was transferred to conical tubes and centrifuged at 2,800 rpm for 13 minutes at 4 °C with a deceleration (DCC) setting of 5. The plasma supernatant was carefully transferred to fresh tubes without disturbing the buffy coat and subjected to a second centrifugation at 3,700 rpm for 13 minutes at 4 °C (DCC 5) to remove residual cellular debris. Plasma was gently mixed, aliquoted into cryovials, and stored at -80 °C until downstream analyses. All samples were coded and de-identified prior to research use.

##### 1.3.3 cfDNA purification

cfDNA purification was performed with Zymogen Quick-cfDNA Serum & Plasma Kit®. 2 mL of plasma from each patient was subjected to column-based processing using the Zymo-Spin™ III-S system. Plasma was lysed with protease K and DNA digestion buffer at 55 °C for 30 minutes, then mixed with DNA binding buffer, loaded onto columns and centrifuged at 1,000 × g for 2 minutes, with repeated spins until the full volume passed through the column. Columns were then washed sequentially with DNA Prep Buffer and DNA Wash Buffer, each followed by centrifugation at ≥10,000 × g for 30 seconds. A final high-speed centrifugation at full speed ( 15,000 × g) for 1 minute ensured complete removal of residual wash buffer. DNA was eluted by incubating columns with 42 µl elution buffer for 3 minutes at room temperature, followed by centrifugation at maximum speed for 30 seconds. 1 µl of elution was taken for concentration measurement with Quibit 4.0 High Sensitivity dsDNA module. Another 1 µl of elution was taken for QC with Bioanalyzer DNA length analysis at UPMC Hillman Cancer Center Genome Facility. All samples used in this study passed QC. Remaining 40 µl purified cfDNA was stored at ≤ -80 °C until downstream analysis.

##### 1.3.4 Sequencing

Ultra-low-pass whole-genome sequencing (ULP-WGS) was performed at the Broad Institute. Library construction was carried out using normalized cfDNA input in the range of 25–52.5 ng in 50 µL of TE buffer (10 mM Tris-HCl, 1 mM EDTA, pH 8.0), based on PicoGreen quantification. Libraries were prepared using the KAPA HyperPrep Kit with Library Amplification (KAPA Biosystems, KK8504) in conjunction with IDT duplex UMI adapters. Unique 8-base dual-index sequences embedded within the p5 and p7 primers (IDT) were added during PCR amplification. Enzymatic clean-up steps were performed using Beckman Coulter AMPure XP beads, with elution volumes reduced to 30 µL to maximize library concentration. Following library preparation, library concentrations were measured using the Invitrogen Quant-IT broad range dsDNA assay kit (Thermo Scientific, Q33130) with a 1:200 PicoGreen dilution. Each library was then normalized to a concentration of 35 ng/µL using 10 mM Tris-HCl (pH 8.0). For ULP-WGS, approximately 4 µL of each normalized library was transferred to a new tube and further diluted to 2 ng/µL using 10 mM Tris-HCl (pH 8.0). Up to 95 ULP-WGS libraries were pooled using equivolume pooling. The final pool was quantified by qPCR and normalized to the appropriate concentration for sequencing. Cluster amplification of pooled libraries was performed according to the manufacturer's protocol (Illumina) using exclusion amplification cluster chemistry on NovaSeq SP flow cells. Sequencing was carried out using sequencing-by-synthesis chemistry on the NovaSeq platform with paired-end 151 bp reads. The target range of coverage is 0.1x to 0.3x, and the final mean coverage of sequencing is 0.21x.

##### 1.3.5 Analysis pipeline for tumor fraction estimation and fragment length

Following sequencing, the Broad Institute provided aligned BAM files generated against the GRCh38 human reference genome. Alignment quality and sequencing performance metrics were assessed using FASTQC (by Babraham Bioinformatics) and Qualimap to ensure ultra-low coverage, mapping quality, and overall data quality [4]. Tumor fraction (TFx) estimation from cfDNA was performed using ichorCNA, which applies a hidden Markov model (HMM) based probabilistic framework to infer the fraction of tumor-derived DNA in plasma. The reference genome, segmented into fixed-width bins (1,000 kb), was used as the optimized resolution. Prior to TFx estimation, GC-content bias was corrected using the HMMcopy, as implemented in the ichorCNA workflow. The GC-corrected and normalized data were then used to estimate the tumor fraction for each sample [5]. To characterize cfDNA fragment size distributions, insert size metrics were computed from the aligned BAM files using Picard (v3.0.0) CollectInsertSizeMetrics [6], providing summary statistics and distributions of fragment lengths for downstream analyses.

[7] Carmen Martin-Alonso, Shervin Tabrizi, Kan Xiong, Timothy Blewett, Sainetra Sridhar, Andjela Crnjac, Sahil Patel,

Zhenyi An, Ahmet Bekdemir, Douglas Shea, et al. Priming agents transiently reduce the clearance of cell-free DNA to improve liquid biopsies. *Science*, 383(6680):eadf2341, 2024.

#### 2 Supplementary Figures

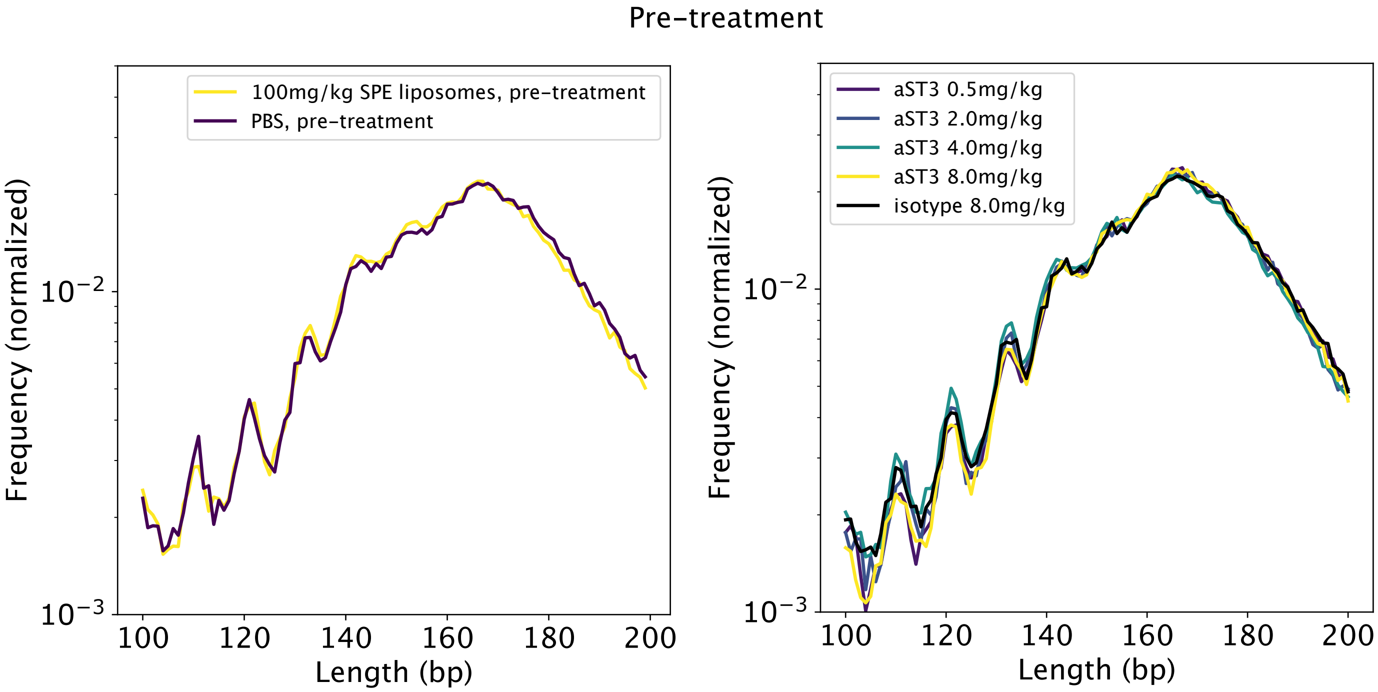

Figure S1: Aggregate fragment distribution plots as in **Figure 2 C-D**, for pre-treatment mice.

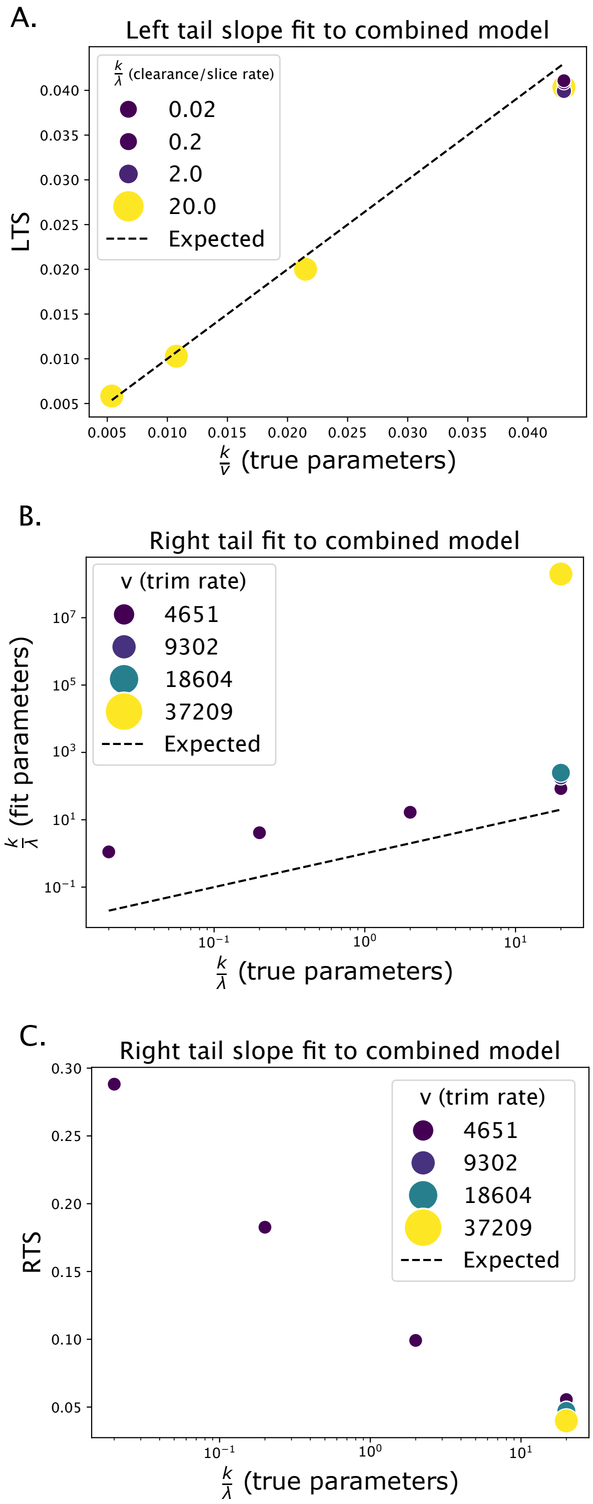

Figure S2: Parameter inference on combined model. **A:** The inferred left tail slope of the simulations from **Figure 2** compared to $\frac{k}{v}$. The dot size corresponds to the value of $\frac{k}{\lambda}$. **B:** The inferred right tail parameter $\frac{k}{\lambda}$ from least-squares minimization on log-transformed $f_{slice}$ compared to $\frac{k}{\lambda}$, with $\beta$ set to the true value of 0.043 c. The dot size corresponds to the value of $v$. **C:** The inferred right tail slope for the same simulations.

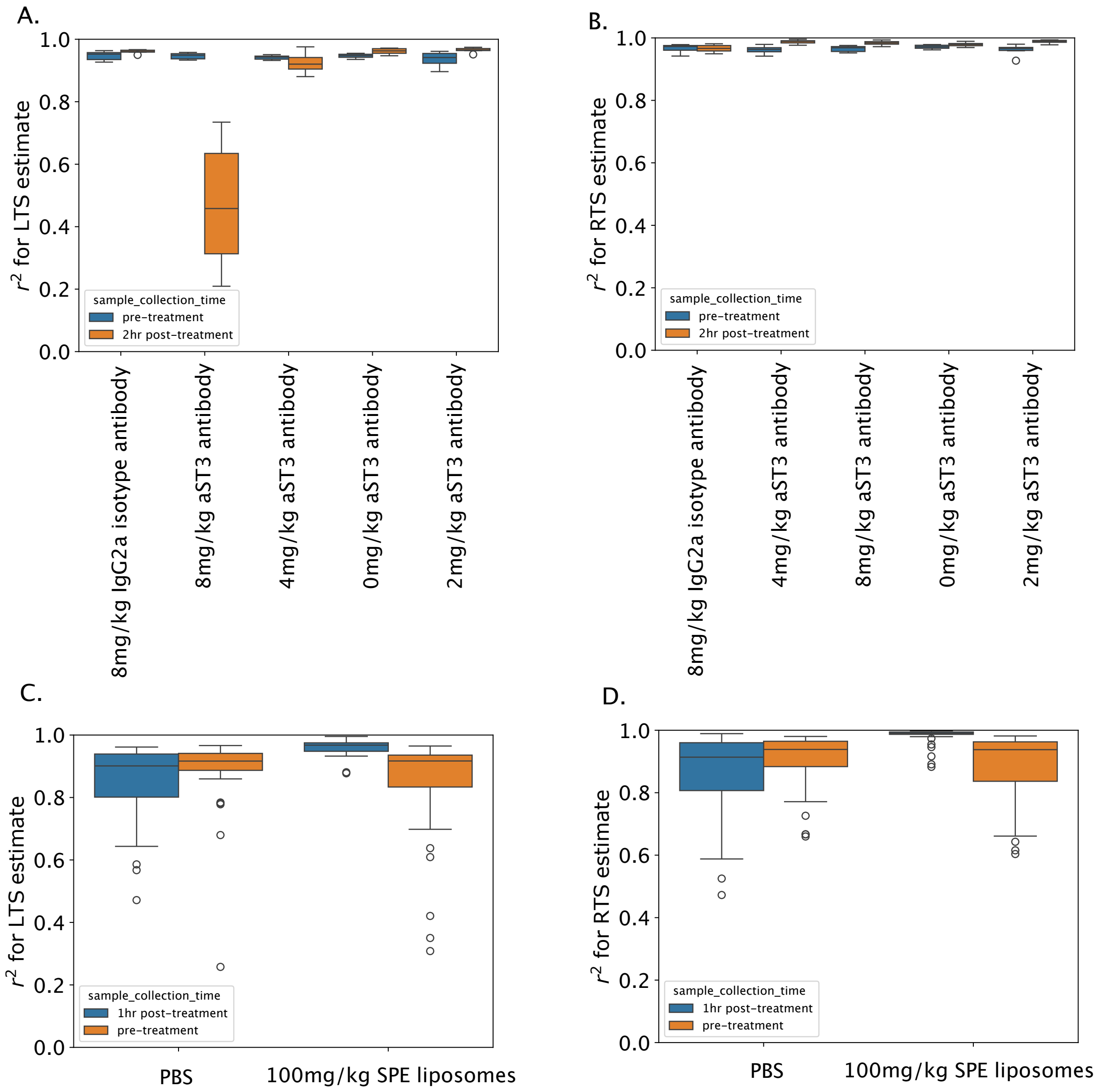

Figure S3: Distribution of Pearson $r^2$ values for the LTS and RTS fit, for all treatments in the MA dataset.

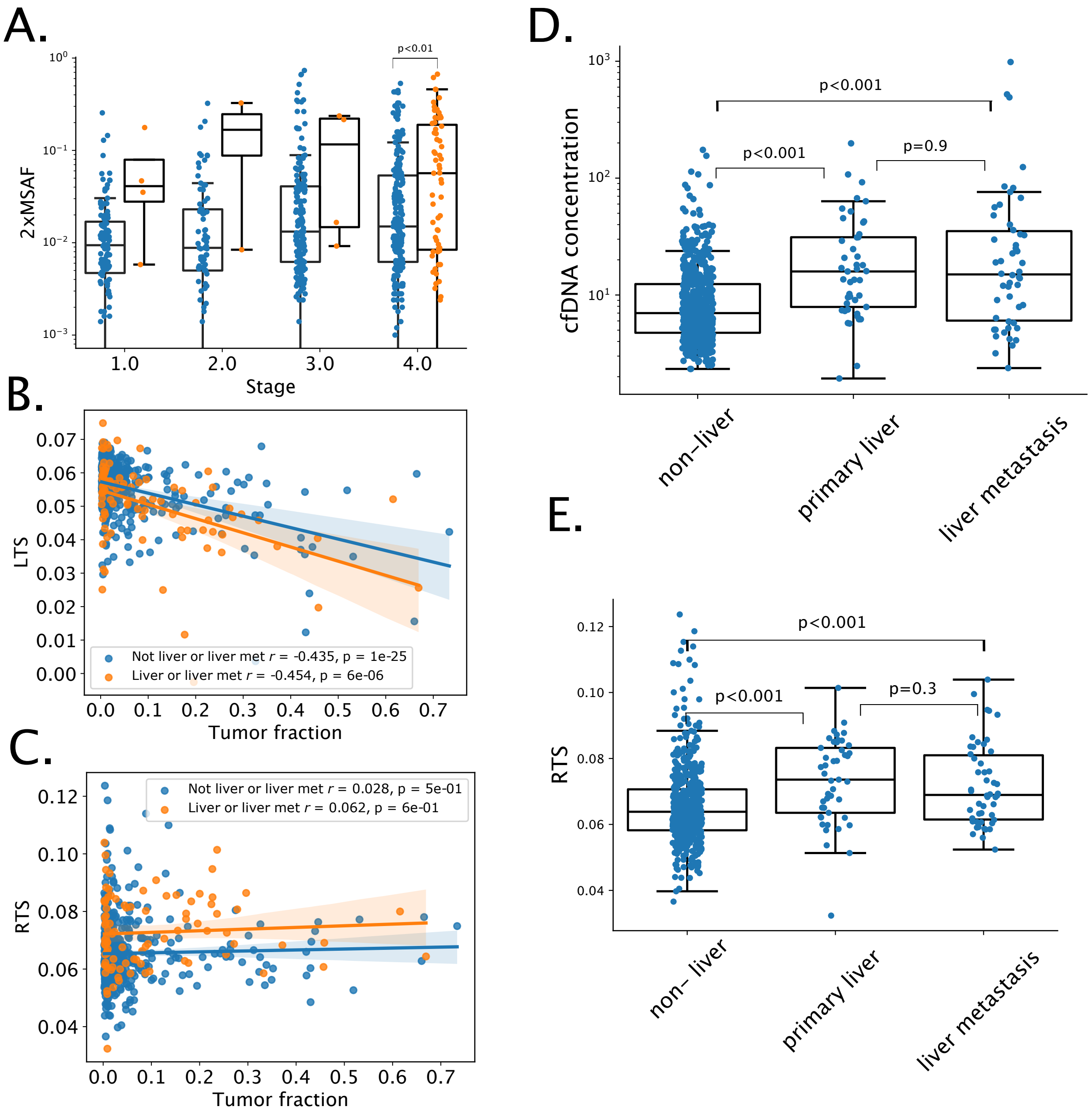

Figure S4: Effects of liver involvement on cfDNA features in Dataset I. **A:** Tumor fraction (2 x MSAF) stratified by Stage and liver involvement for patients in Dataset I. P-values are computed from a pairwise Mann-Whitney U-test and are BH-corrected with a FDR of 5% for between-stage comparisons. For complete statistics see **Tables S5, S6, S8**. **B-C:** Relationship between tumor fraction, LTS and RTS as in **Main Figure 3** stratifying by liver involvement. **D-E:** Comparison of liver involvement to non-involvement controlling for primary liver cancer or liver metastasis. P-values are corrected via BH with a FDR of 5%.

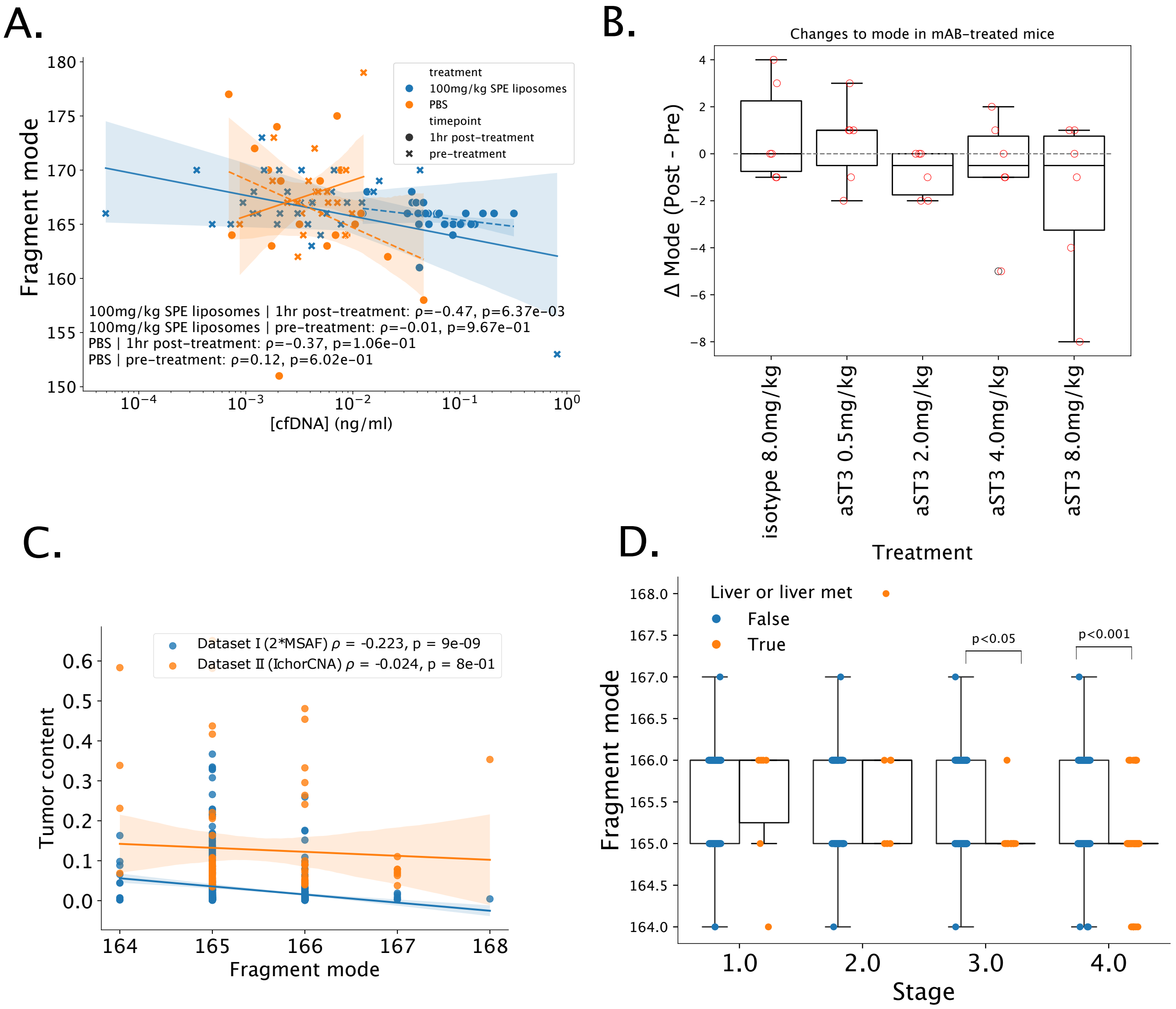

Figure S5: Supporting plots for mode fragment length results. **A:** Correlation between fragment mode and cfDNA concentration in the nanoparticle treatment. **B:** Change in mode plotted against antibody treatment. **C:** Relationship between mode fragment length and tumor content for both cancer datasets. **D:** Mode fragment length stratified by Stage and liver involvement for patients in Dataset I. P-values from a pairwise Mann-Whitney U-test are Bonferroni-corrected for between-stage comparisons. For complete statistics see **Table S7**.

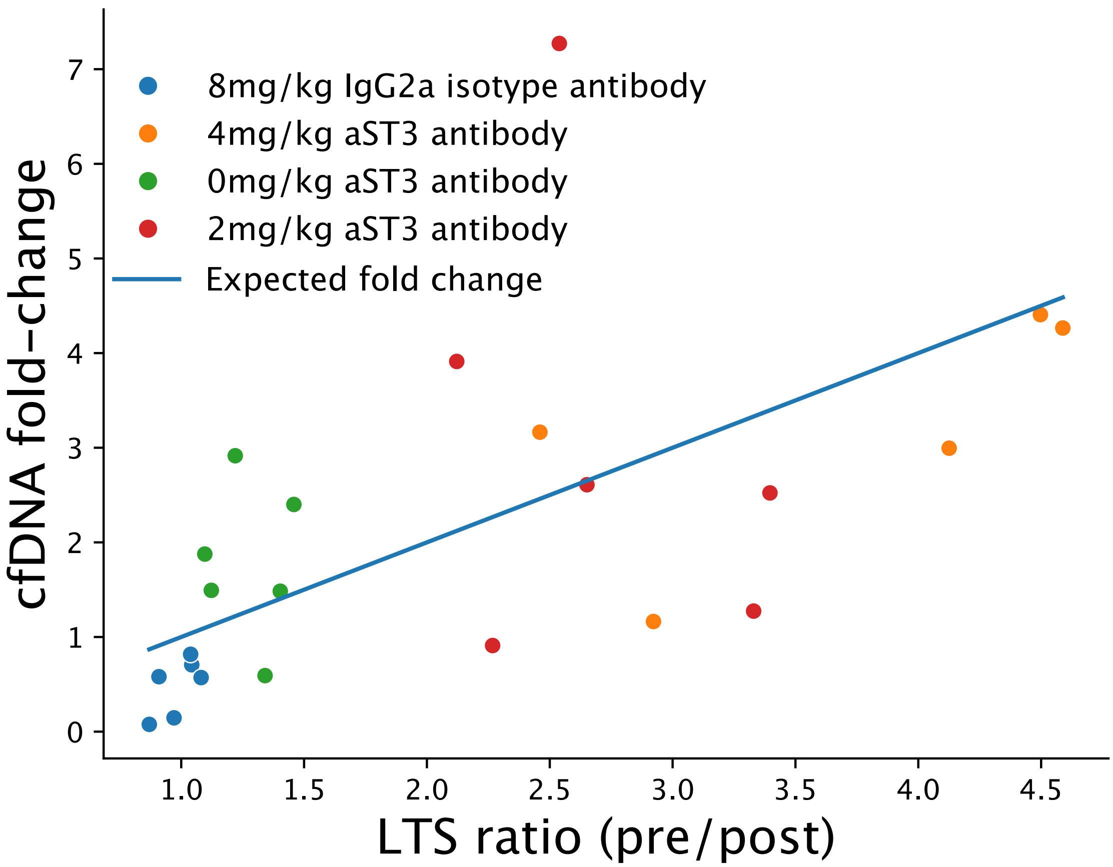

Figure S6: Relationship between cfDNA fold change and LTS ratio in antibody-treated mice. The 8mg/kg aST3 treatment is not shown because of a poor LTS fit (median $r^2$ of 0.5). Points indicate the dosage and the blue line corresponds to y=x. The LTS ratio $\frac{LTS_{pre}}{LTS_{post}}$ is $\frac{k_{pre}}{v_{pre}} \cdot \frac{v_{post}}{k_{post}}$. If $v_{pre} = v_{post}$, so that circulation time rather than absolute fragmentation rate determines LTS, then $\frac{LTS_{pre}}{LTS_{post}} = \frac{k_{pre}}{k_{post}}$. If [cfDNA] $\propto \frac{1}{k}$ then $\frac{LTS_{pre}}{LTS_{post}} \approx \frac{\text{cfDNA}_{post}}{\text{cfDNA}_{pre}}$.

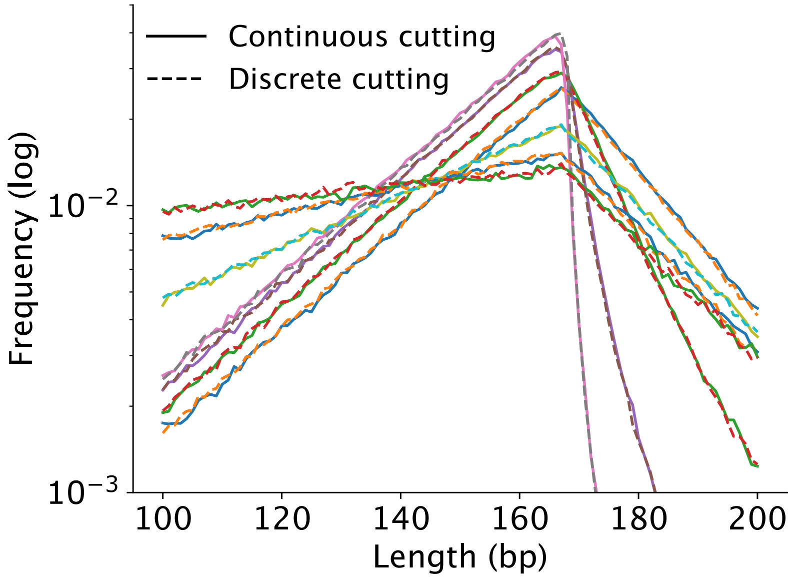

Figure S7: Combined model simulations with identical parameters to **Main Figure 2A-B** for both a model where linker regions are cut continuously along the fragment and binned into integers at the end of the simulation (solid), or cut at discrete basepairs during the simulation (dashed).

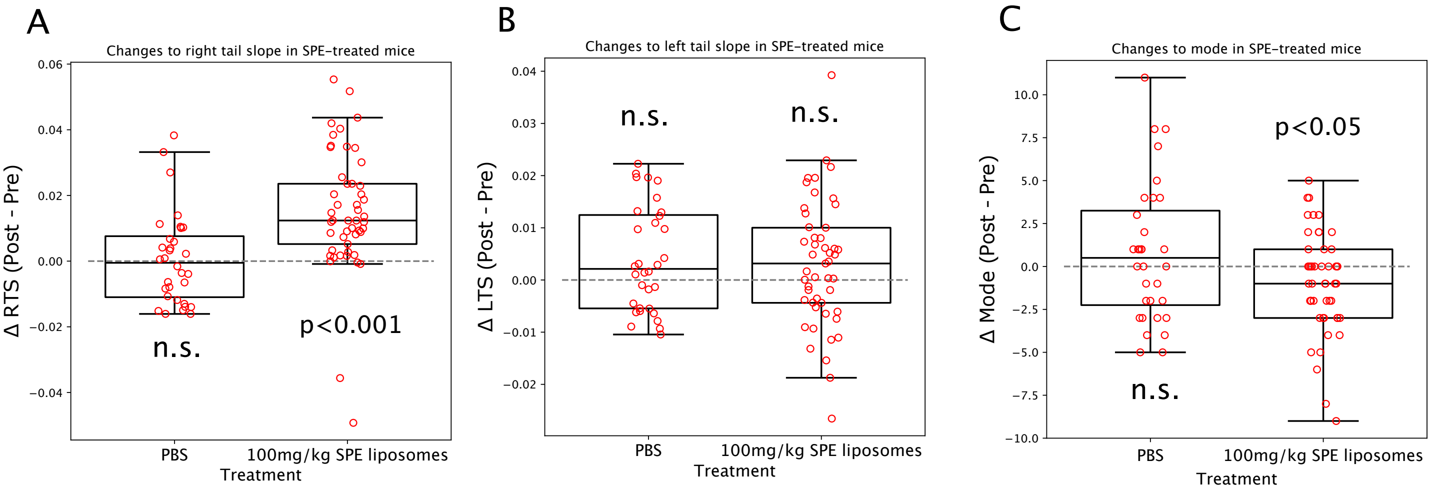

Figure S8: Fragment length comparisons using paired retro-orbital samples only (PBS n=32, Nanoparticle n=49). **A-C:** Pairwise changes in RTS, LTS, and fragment mode respectively with p-values from the two-sided one-sample Wilcoxon signed rank test ($\alpha = 0.05$).

### 3 Supplementary Tables

Table S1: Dataset I characteristics

| Variable | Category | N | Percent | Mean ± SD | Q1 | Q2 | Q3 | Q4 |
|---|---|---|---|---|---|---|---|---|
| Age (years) | | 696 | 100.0 | 64.3 ± 13.1 | 57.0 | 66.0 | 74.0 | 99 |
| PFS (months) | | 683 | 98.1 | 15.65 ± 11.86 | 5.20 | 12.73 | 25.32 | 41.47 |
| | NA | 13 | 1.9 | | | | | |
| DSS (months) | | 685 | 98.4 | 19.69 ± 11.64 | 10.20 | 17.63 | 30.37 | 41.47 |
| | NA | 11 | 1.6 | | | | | |
| Plasma cfDNA concentration (ng/ml) | | 696 | 100.0 | 16.24 ± 48.94 | 5.00 | 7.45 | 14.40 | 985.00 |
| Tumor fraction (MSAF) | | 696 | 100.0 | 0.025 ± 0.050 | 0.003 | 0.006 | 0.021 | 0.367 |
| Sex | Female | 357 | 51.3 | | | | | |
| | Male | 339 | 48.7 | | | | | |
| Intent | Adjuvant | 52 | 7.5 | | | | | |
| | Curative | 302 | 43.4 | | | | | |
| | Palliative | 326 | 46.8 | | | | | |
| | Surveillance | 4 | 0.6 | | | | | |
| | NA | 12 | 1.7 | | | | | |
| Stage | 0.0 | 2 | 0.3 | | | | | |
| | 1.0 | 114 | 16.3 | | | | | |
| | 2.0 | 78 | 11.2 | | | | | |
| | 3.0 | 173 | 24.9 | | | | | |
| | 4.0 | 300 | 43.1 | | | | | |
| | NA | 29 | 4.2 | | | | | |
| Progressed | No | 333 | 47.8 | | | | | |
| | Yes | 350 | 50.3 | | | | | |
| | NA | 13 | 1.9 | | | | | |
| Deceased | No | 441 | 63.4 | | | | | |
| | Yes | 244 | 35.1 | | | | | |
| | NA | 11 | 1.6 | | | | | |

Table S2: Dataset I stages by tumor type

| Stage<br>Tumor type | Total | 0 | 1 | 2 | 3 | 4 | NA |
|---|---|---|---|---|---|---|---|
| Appendix | 61 | 0 | 2 | 3 | 1 | 55 | 0 |
| Bladder/Urothelial | 23 | 0 | 2 | 8 | 2 | 10 | 1 |
| Blood | 20 | 0 | 3 | 4 | 6 | 5 | 2 |
| Brain | 15 | 0 | 2 | 1 | 0 | 2 | 10 |
| Breast | 43 | 0 | 10 | 11 | 14 | 8 | 0 |
| Cervix | 16 | 1 | 6 | 0 | 3 | 6 | 0 |
| Colon/Rectum | 43 | 0 | 1 | 3 | 7 | 32 | 0 |
| Endocrine | 6 | 0 | 3 | 1 | 0 | 2 | 0 |
| Endometrium (uterus) | 45 | 0 | 15 | 2 | 7 | 17 | 4 |
| Esophagus/Pharynx | 25 | 0 | 1 | 2 | 8 | 14 | 0 |
| Head/neck | 25 | 0 | 5 | 4 | 11 | 5 | 0 |
| Hepatobiliary | 37 | 0 | 3 | 7 | 12 | 15 | 0 |
| Kidney | 28 | 0 | 9 | 2 | 3 | 13 | 1 |
| Liver | 9 | 0 | 3 | 1 | 2 | 3 | 0 |
| Lung | 70 | 0 | 7 | 8 | 23 | 32 | 0 |
| Ovary/Fallopian Tubes | 36 | 0 | 4 | 1 | 15 | 16 | 0 |
| Pancreas | 23 | 0 | 6 | 3 | 4 | 10 | 0 |
| Prostate | 13 | 0 | 0 | 0 | 2 | 11 | 0 |
| Skin | 46 | 0 | 14 | 6 | 14 | 12 | 0 |
| Small intestine | 16 | 0 | 2 | 0 | 7 | 7 | 0 |
| Soft Tissue | 63 | 0 | 8 | 7 | 21 | 18 | 9 |
| Stomach | 20 | 0 | 6 | 3 | 6 | 5 | 0 |
| Testicle | 7 | 0 | 0 | 1 | 5 | 0 | 1 |
| Unknown | 2 | 0 | 0 | 0 | 0 | 2 | 0 |
| Vagina/Vulva | 2 | 0 | 2 | 0 | 0 | 0 | 0 |

Table S3: Dataset II cfDNA and tumor fraction

| **Patient Index** | **Samples** | **Plasma cfDNA concentration (ng/ml)** | | | | | | | **IchorCNA tumor fraction (%)** | | | | | | |
|---|---|---|---|---|---|---|---|---|---|---|---|---|---|---|---|
| | | mean | std | min | 25% | 50% | 75% | max | mean | std | min | 25% | 50% | 75% | max |
| 0 | 1 | 19.4 | | 19.4 | 19.4 | 19.4 | 19.4 | 19.4 | 6.01 | | 6.01 | 6.01 | 6.01 | 6.01 | 6.01 |
| 1 | 5 | 27.9 | 16.4 | 12.3 | 20.6 | 22.4 | 29 | 55.2 | 7.64 | 1.8 | 6.15 | 6.47 | 6.98 | 7.98 | 10.6 |
| 2 | 4 | 10.5 | 3.91 | 7.88 | 8.06 | 8.88 | 11.3 | 16.2 | 8.19 | 7 | 0 | 4.21 | 8.2 | 12.2 | 16.3 |
| 3 | 6 | 42.2 | 25.1 | 15.8 | 25.6 | 33.9 | 59.2 | 78.8 | 5 | 3.39 | 0 | 3.67 | 4.48 | 7.17 | 9.54 |
| 4 | 2 | 9.14 | 0.537 | 8.76 | 8.95 | 9.14 | 9.33 | 9.52 | 3.93 | 5.56 | 0 | 1.97 | 3.93 | 5.9 | 7.87 |
| 5 | 2 | 68.9 | 52.5 | 31.8 | 50.4 | 68.9 | 87.5 | 106 | 15.3 | 11.1 | 7.4 | 11.3 | 15.3 | 19.2 | 23.1 |
| 6 | 6 | 11.7 | 3.52 | 8.64 | 10.2 | 10.6 | 11.5 | 18.6 | 6.83 | 2.61 | 3.76 | 4.55 | 7.06 | 8.8 | 9.98 |
| 7 | 2 | 9.5 | 0.877 | 8.88 | 9.19 | 9.5 | 9.81 | 10.1 | 2.74 | 3.87 | 0 | 1.37 | 2.74 | 4.1 | 5.47 |
| 8 | 3 | 31.6 | 11.4 | 21.6 | 25.4 | 29.2 | 36.6 | 44 | 6.16 | 1.71 | 4.57 | 5.25 | 5.94 | 6.95 | 7.96 |
| 9 | 5 | 11.4 | 0.874 | 10 | 11.4 | 11.6 | 12 | 12.2 | 10.1 | 6.28 | 5.42 | 6.62 | 8.51 | 8.75 | 21 |
| 10 | 4 | 26 | 6.15 | 21.2 | 22.1 | 24.1 | 28.1 | 34.8 | 7.2 | 1.81 | 4.87 | 6.29 | 7.49 | 8.41 | 8.96 |
| 11 | 5 | 28.5 | 28.1 | 4.72 | 10.2 | 22.4 | 29.6 | 75.6 | 9.75 | 7.07 | 5 | 5.53 | 7.04 | 9.09 | 22.1 |
| 12 | 2 | 20.5 | 12.5 | 11.7 | 16.1 | 20.5 | 25 | 29.4 | 7.9 | 1.47 | 6.86 | 7.38 | 7.9 | 8.42 | 8.94 |
| 13 | 2 | 16.7 | 10.6 | 9.28 | 13 | 16.7 | 20.5 | 24.2 | 6.03 | 2.42 | 4.32 | 5.18 | 6.03 | 6.89 | 7.75 |
| 14 | 4 | 7.19 | 3.99 | 3.04 | 5.53 | 6.54 | 8.2 | 12.6 | 30.3 | 14.4 | 11 | 24.9 | 32.4 | 37.9 | 45.4 |
| 15 | 5 | 9.2 | 4.31 | 4.48 | 5.56 | 9.48 | 11.5 | 15 | 4.44 | 2.61 | 0 | 4.66 | 5.03 | 5.72 | 6.79 |
| 16 | 3 | 26.5 | 19.9 | 10.6 | 15.4 | 20.2 | 34.5 | 48.8 | 2.66 | 2.52 | 0 | 1.47 | 2.95 | 3.98 | 5.02 |
| 17 | 4 | 150 | 147 | 34.8 | 79.2 | 100 | 171 | 366 | 30.9 | 19.5 | 5.26 | 21.1 | 35 | 44.8 | 48.1 |
| 18 | 2 | 13.5 | 12.1 | 4.92 | 9.19 | 13.5 | 17.7 | 22 | 7.91 | 1.02 | 7.18 | 7.54 | 7.91 | 8.27 | 8.63 |
| 19 | 2 | 20.3 | 5.23 | 16.6 | 18.5 | 20.3 | 22.1 | 24 | 7.18 | 3.7 | 4.56 | 5.87 | 7.18 | 8.49 | 9.8 |
| 20 | 1 | 18 | | 18 | 18 | 18 | 18 | 18 | 6.89 | | 6.89 | 6.89 | 6.89 | 6.89 | 6.89 |
| 21 | 2 | 6.62 | 1.16 | 5.8 | 6.21 | 6.62 | 7.03 | 7.44 | 4.04 | 5.71 | 0 | 2.02 | 4.04 | 6.06 | 8.07 |
| 22 | 2 | 44.3 | 6.93 | 39.4 | 41.9 | 44.3 | 46.8 | 49.2 | 21 | 17.4 | 8.68 | 14.8 | 21 | 27.1 | 33.2 |
| 23 | 3 | 13.4 | 9.89 | 7.08 | 7.7 | 8.32 | 16.6 | 24.8 | 23.3 | 18.8 | 4.08 | 14.1 | 24.1 | 32.9 | 41.7 |
| 24 | 3 | 15.3 | 6.73 | 9 | 11.8 | 14.6 | 18.5 | 22.4 | 5.45 | 1.58 | 4.15 | 4.58 | 5 | 6.11 | 7.21 |
| 25 | 2 | 28 | 2.83 | 26 | 27 | 28 | 29 | 30 | 4.27 | 0.54 | 3.89 | 4.08 | 4.27 | 4.46 | 4.65 |
| 26 | 2 | 7.42 | 0.933 | 6.76 | 7.09 | 7.42 | 7.75 | 8.08 | 6.46 | 1.86 | 5.14 | 5.8 | 6.46 | 7.11 | 7.77 |
| 27 | 3 | 14.5 | 2.67 | 12.9 | 12.9 | 13 | 15.3 | 17.6 | 4.3 | 3.79 | 0 | 2.88 | 5.75 | 6.45 | 7.16 |
| 28 | 3 | 90.6 | 109 | 17 | 27.9 | 38.8 | 127 | 216 | 19.7 | 14.6 | 4.63 | 12.6 | 20.5 | 27.2 | 33.9 |
| 29 | 2 | 7.7 | 2.23 | 6.12 | 6.91 | 7.7 | 8.49 | 9.28 | 3.43 | 0.622 | 2.99 | 3.21 | 3.43 | 3.65 | 3.87 |
| 30 | 3 | 218 | 66.6 | 148 | 187 | 226 | 253 | 280 | 60.5 | 3.97 | 58.1 | 58.2 | 58.3 | 61.7 | 65.1 |
| overall | 95 | 34.3 | 57 | 3.04 | 9.58 | 16.6 | 29.5 | 366 | 11.5 | 13.6 | 0 | 4.66 | 6.89 | 9.33 | 65.1 |

Table S4: Key parameters of the cfDNA fragmentation-clearance simulation.

| Symbol | Code name | Description | Default value | Units |
|---|---|---|---|---|
| $s$ | `s` | Entry (birth) rate of new fragments. | $10^8$ | fragments/time |
| $\lambda_{\text{linker}}$ | `lam_linker` | Per-base-pair linker fragmentation rate; total fragmentation propensity is $\lambda_{\text{linker}} \sum_i (L_{\text{left},i} + L_{\text{right},i})$. | 10 | $(\text{bp}\cdot\text{time})^{-1}$ |
| $v$ | `v_end` | External cut rate per fragment. | 4651 | $\text{time}^{-1}$ |
| $k$ | `k` | Clearance (exit) rate per fragment. | 200 | $\text{time}^{-1}$ |
| $n_0$ | `n0` | Initial nucleosome length on entry; nucleosome is eroded by end trimming. | 167 | bp |
| $\ell_0$ | `l0` | Baseline linker length added on entry (before exponential noise). | 0 | bp |
| $\beta$ | `b` | Rate parameter for entry linker noise: $X \sim \text{Exponential}(\text{rate} = \beta)$, so $L_0 = \ell_0 + X$. | 0.043 | bp |
| $t_{\text{max}}$ | `t_max` | Maximum simulation time | 100 | a.u. |
| $N_{\text{max}}$ | `max_events` | Maximum number of Gillespie events | $10^8$ | events |

Table S5: Pairwise tests for cfDNA stratified by stage and liver involvement.

| Contrast | Stage | A | B | U-val | alternative | p-unc | p-corr |
|---|---|---|---|---|---|---|---|
| Stage | - | 1 | 2 | 4.87e+03 | two-sided | 0.267 | 0.267 |
| Stage | - | 1 | 3 | 8.8e+03 | two-sided | 0.165 | 0.198 |
| Stage | - | 1 | 4 | 1.28e+04 | two-sided | 9.88e-05 | 0.000296 |
| Stage | - | 2 | 3 | 5.36e+03 | two-sided | 0.0133 | 0.02 |
| Stage | - | 2 | 4 | 7.63e+03 | two-sided | 2.93e-06 | 1.76e-05 |
| Stage | - | 3 | 4 | 2.12e+04 | two-sided | 0.0026 | 0.00519 |
| Liver or liver met | - | False | True | 1.54e+04 | two-sided | 2.71e-11 | |
| Stage * Liver or liver met | 1 | False | True | 209 | two-sided | 0.146 | 0.195 |
| Stage * Liver or liver met | 2 | False | True | 210 | two-sided | 0.256 | 0.256 |
| Stage * Liver or liver met | 3 | False | True | 394 | two-sided | 0.000225 | 0.00045 |
| Stage * Liver or liver met | 4 | False | True | 5e+03 | two-sided | 6.5e-06 | 2.6e-05 |

Table S6: Pairwise tests for RTS stratified by stage and liver involvement.

| Contrast | Stage | A | B | U-val | alternative | p-unc | p-corr |
|---|---|---|---|---|---|---|---|
| Stage | - | 1 | 2 | 4.7e+03 | two-sided | 0.494 | 0.494 |
| Stage | - | 1 | 3 | 8.6e+03 | two-sided | 0.0917 | 0.11 |
| Stage | - | 1 | 4 | 1.19e+04 | two-sided | 3e-06 | 1.8e-05 |
| Stage | - | 2 | 3 | 5.58e+03 | two-sided | 0.0389 | 0.0584 |
| Stage | - | 2 | 4 | 7.78e+03 | two-sided | 7.14e-06 | 2.14e-05 |
| Stage | - | 3 | 4 | 2.08e+04 | two-sided | 0.000877 | 0.00175 |
| Liver or liver met | - | False | True | 1.77e+04 | two-sided | 8.32e-08 | |
| Stage * Liver or liver met | 1 | False | True | 293 | two-sided | 0.706 | 0.706 |
| Stage * Liver or liver met | 2 | False | True | 310 | two-sided | 0.633 | 0.706 |
| Stage * Liver or liver met | 3 | False | True | 432 | two-sided | 0.000531 | 0.00106 |
| Stage * Liver or liver met | 4 | False | True | 5.46e+03 | two-sided | 0.000155 | 0.00062 |

Table S7: Pairwise tests for fragment mode stratified by stage and liver involvement.

| Contrast | Stage | A | B | U-val | alternative | p-unc | p-corr |
|---|---|---|---|---|---|---|---|
| Stage | - | 1 | 2 | 4.59e+03 | two-sided | 0.668 | 0.668 |
| Stage | - | 1 | 3 | 1.11e+04 | two-sided | 0.025 | 0.0375 |
| Stage | - | 1 | 4 | 2.19e+04 | two-sided | 1.43e-07 | 8.6e-07 |
| Stage | - | 2 | 3 | 7.36e+03 | two-sided | 0.134 | 0.16 |
| Stage | - | 2 | 4 | 1.46e+04 | two-sided | 4.12e-05 | 0.000123 |
| Stage | - | 3 | 4 | 2.95e+04 | two-sided | 0.000711 | 0.00142 |
| Liver or liver met | - | False | True | 3.45e+04 | two-sided | 2.95e-07 | |
| Stage * Liver or liver met | 1 | False | True | 320 | two-sided | 0.953 | 0.953 |
| Stage * Liver or liver met | 2 | False | True | 235 | two-sided | 0.405 | 0.54 |
| Stage * Liver or liver met | 3 | False | True | 1.42e+03 | two-sided | 0.0076 | 0.0152 |
| Stage * Liver or liver met | 4 | False | True | 9.95e+03 | two-sided | 3.09e-05 | 0.000124 |

Table S8: Pairwise tests for tumor fraction (2xMSAF) stratified by stage and liver involvement.

| Contrast | Stage | A | B | U-val | alternative | p-unc | p-corr |
|---|---|---|---|---|---|---|---|
| Stage | - | 1 | 2 | 4.32e+03 | two-sided | 0.74 | 0.74 |
| Stage | - | 1 | 3 | 7.45e+03 | two-sided | 0.000745 | 0.00149 |
| Stage | - | 1 | 4 | 1.2e+04 | two-sided | 4.49e-06 | 2.69e-05 |
| Stage | - | 2 | 3 | 5.38e+03 | two-sided | 0.0145 | 0.0217 |
| Stage | - | 2 | 4 | 8.64e+03 | two-sided | 0.000486 | 0.00146 |
| Stage | - | 3 | 4 | 2.37e+04 | two-sided | 0.198 | 0.238 |
| Liver or liver met | - | False | True | 2.07e+04 | two-sided | 0.000328 | |
| Stage * Liver or liver met | 1 | False | True | 229 | two-sided | 0.23 | 0.461 |
| Stage * Liver or liver met | 2 | False | True | 293 | two-sided | 0.837 | 0.837 |
| Stage * Liver or liver met | 3 | False | True | 962 | two-sided | 0.707 | 0.837 |
| Stage * Liver or liver met | 4 | False | True | 5.97e+03 | two-sided | 0.00312 | 0.0125 |

Table S9: Counts for samples used from [7] after filtering

| Treatment | Sample collection time | Sample collection type | $n$ |
|---|---|---|---|
| 100mg/kg SPE liposomes | 1hr post-treatment | retro-orbital | 52 |
| 100mg/kg SPE liposomes | pre-treatment | retro-orbital | 51 |
| PBS | pre-treatment | retro-orbital | 35 |
| PBS | 1hr post-treatment | retro-orbital | 34 |
| 100mg/kg SPE liposomes | 1hr post-treatment | terminal bleed | 16 |
| PBS | 1hr post-treatment | terminal bleed | 12 |
| 0.5mg/kg aST3 antibody | 2hr post-treatment | terminal bleed | 6 |
| 0.5mg/kg aST3 antibody | pre-treatment | retro-orbital | 6 |
| 2mg/kg aST3 antibody | 2hr post-treatment | terminal bleed | 6 |
| 2mg/kg aST3 antibody | pre-treatment | retro-orbital | 6 |
| 4mg/kg aST3 antibody | 2hr post-treatment | terminal bleed | 6 |
| 4mg/kg aST3 antibody | pre-treatment | retro-orbital | 6 |
| 8mg/kg IgG2a isotype antibody | 2hr post-treatment | terminal bleed | 6 |
| 8mg/kg IgG2a isotype antibody | pre-treatment | retro-orbital | 6 |
| 8mg/kg aST3 antibody | 2hr post-treatment | terminal bleed | 6 |
| 8mg/kg aST3 antibody | pre-treatment | retro-orbital | 6 |
